# Seabird diet and transmission routes mediate the increase in pathogen prevalence with human disturbance

**DOI:** 10.64898/2026.01.08.698262

**Authors:** Shabnam Shadloo, Amy Wilson, Marie Auger-Méthé

## Abstract

Human disturbances can alter patterns of wildlife infection, but their effects are likely to differ between terrestrial and marine environments. Seabirds range across land and sea and are therefore exposed to diverse human pressures such as urbanization, agriculture, and landfills, as well as fisheries, shipping, and marine pollution. However, most literature on wildlife infection has focused on terrestrial hosts or single type of disturbance (e.g., urbanization). Studying seabirds, which exploit diverse diets and habitats, offers a unique opportunity to disentangle how disturbances at sea and on land, as well as diets, and transmission routes, can change pathogen prevalence at a global scale. We conducted a global literature review and extracted datasets on seabird infection status for a range of pathogens, seabirds diets (piscivore, invertivore, generalist), and pathogen transmission routes (contact-based, vector-borne, intermediate host) to link pathogen prevalence to human disturbances at sea and on land. Our results showed that the association between human disturbances and pathogen prevalence was mediated by species’ diets and differed across transmission routes. Across gradients of human disturbance at sea, pathogen prevalence in piscivorous seabirds increased from near zero to over 0.7, particularly for contact-based pathogens. Land-based disturbance was also associated with higher pathogen prevalence in generalists across transmission routes, whereas invertivores showed a negative association with terrestrial disturbance. These results indicate that piscivores are most vulnerable in heavily disturbed marine environments, while generalists face greater risk on highly disturbed lands, suggesting that effective disease mitigation will require targeted management actions in both environments.

## 2 Introduction

Disease outbreaks in wildlife occur through intrinsic and extrinsic factors and can be accelerated by human activities (Daszak, Cunningham, and Hyatt, 2000; Bradley and Altizer, 2007). These outbreaks pose serious threats to species by leading to higher mortality, reduced reproductive success, population decline, local extinction, and biodiversity loss (Brearley et al., 2013; Khan et al., 2019; Sanz-Aguilar et al., 2020; Camphuysen, Gear, and Furness, 2022)). For example, viral and bacterial diseases, including avian cholera (*Pasteurella multocida*), have caused significant mortality in aquatic birds and can lead to rapid population decline and even local extinction (Newman et al., 2007; Iverson et al., 2016; Descamps et al., 2012). Recently, highly pathogenic avian influenza panzootic decimated seabird populations in North and South America, Europe, and Antarctica (McPhail et al., 2024; Grémillet et al., 2023). Over three billion people live in areas with frequent contact with wildlife, further increasing the risk of disease transmission (Simkin et al., 2025). Urbanization creates opportunities for interactions between wildlife, livestock, and humans, modifying pathogen transmission through host density, movement, and immunity, as well as abiotic factors such as pollution and climate change (Hassell et al., 2017). Seabirds are especially vulnerable wildlife species. They are effective reservoirs for pathogens as they live long, breed in dense colonies, and have diverse diets. They also travel long distances, sometimes across continents, crossing different gradients of human impact (Khan et al., 2019; Dietrich, Gómez-Díaz, and McCoy, 2011; McCoy et al., 2016; Dhama, Mahendran, and Tomar, 2008). Half of seabird species are listed as globally threatened or near threatened by the IUCN (Dias et al., 2019). This situation may worsen due to human-related factors such as climate change, pollution, and the dilution effect due to loss of biodiversity (Cohen et al., 2020; Grimaldi et al., 2015; Keesing et al., 2010; Bradley and Altizer, 2007).

Anthropogenic factors on terrestrial environments, such as urban development, alter seabirds’ environments and behaviour, influencing their interactions with pathogens (Matos, Domit, and Bracarense, 2020; Hassell et al., 2017; Morse et al., 2012). Human activities concentrate food resources and pathogen hosts in new ways. For instance, open-air landfills and agricultural sites provide abundant, predictable food that attracts large numbers of opportunistic seabirds. However, these dense foraging aggregations become hotspots for pathogen transmission, since many diseases often rely on the spatial overlap of hosts and vectors (Carlberg et al., 2022; Sadeghi et al., 2022; López-Calderón et al., 2023; Hassell et al., 2017; Altizer, Bartel, and Han, 2011; Plowright et al., 2011; Bradley and Altizer, 2007). For example, birds in urbanized landscapes experience higher exposure to mosquito-borne viruses such as West Nile virus (Bradley, Gibbs, and Altizer, 2008). We predict that higher human pressure on terrestrial environments could lead to increased prevalence of pathogens, due to more overlap with intermediate hosts and vectors, and aggregation of species in anthropogenic food sources and contaminated habitats.

Human pressures on marine environments can have a similar impact on seabirds. Marine pollution can introduce new pathogens to seabirds. For example, seabirds foraging near sewage outfalls or wastewater treatment plants often ingest pathogens and antimicrobial-resistant microbes originating from human waste (Hubálek, 2021). Another example is the spread of *Toxoplasma gondii* to marine environments through urban runoffs, leading to infection in seabirds (Poulle et al., 2021). Fishing operations produce abundant waste that can be opportunistically exploited by seabirds (Bartumeus et al., 2010; Cianchetti-Benedetti et al., 2018). Scavenging seabirds congregate in large numbers around fishing vessels or fish processing outfalls and show a strong tendency to repeatedly return to the same feeding locations (Bartumeus et al., 2010). This aggregation around fishing vessels and processing facilities can increase interspecies contact and pathogen exchange. Bycatch can also indirectly increase susceptibility to novel pathogens, especially when it leads to large population declines associated with reduced genetic diversity and resilience (Cortés-Hinojosa, 2024). Many pelagic and polar seabirds may rarely visit cities, but can be affected by marine-based impacts such as bioaccumulation of contaminants and climate-driven shifts in marine pathogens. For example, penguins are exposed to persistent organic pollutants, which can impair their immune function (Jara-Carrasco et al., 2015).

Many offshore seabirds, such as petrels and shearwaters, chronically ingest microplastics, which can increase the prevalence of pathogenic bacteria (Fackelmann et al., 2023; Jersey et al., 2025). Climate change is also shifting the distribution of some pathogens towards higher altitudes, mainly threatening naive species (McCoy et al., 2023; Van Hemert, Pearce, and Handel, 2014; Bradley et al., 2005). Accordingly, we predict that pathogens would occur more in seabirds in waters with higher cumulative human disturbance.

Seabirds’ foraging ecology can also shape how human activities influence the prevalence of pathogens that seabirds encounter. Their foraging ecology can determine movement and habitat use, drawing some species into cities where they access abundant food, while leading others to avoid urban environments. Generalist seabirds are more likely to be exposed to a broader array of pathogens, particularly in anthropogenic landscapes (Gutiérrez et al., 2017). For example, colonies of European herring gulls (*Larus argentatus*) foraging across varied habitats exhibited greater microbial pathogen diversity than those with more specialized diets (Fuirst et al., 2018). However, dietary specialization does not eliminate the risk of pathogens. Foodborne pathogens such as Vibrio spp. can spread through prey species, such as shellfish, fish, zooplankton, and crustaceans (Fuirst et al., 2018; Holzman, 2007). We predict that human-related factors would interact with seabird diet to alter the pathogen prevalence.

The transmission routes used by pathogens can determine which factors accelerate their transmission and how human activities influence their prevalence in seabirds. Transmission occurs through vertical (passed directly from parent to offspring) and/or horizontal routes (Carr and Bronze, 2023). For pathogens to transmit through horizontal routes, suitable ecological and environmental conditions are required, such as direct contact with infected individuals, contaminated environments, or the presence of vectors or intermediate hosts (Pedersen et al., 2005; Thrusfield, 2013). Because human disturbances can modify these conditions, for example, by concentrating animals at landfills, altering prey communities, increasing pollution, and altering food and water-borne pathogens dynamics, the potential effects of humans on pathogen prevalence depend on the transmission route.

Numerous studies have examined pathogen prevalence in wildlife species. Much of this research has focused on the relationship between pathogens and wild mammals (Werner and Nunn, 2020; Albery et al., 2022), species’ habitat use (Gutiérrez et al., 2017)), or has concentrated on specific groups of pathogens (Quillfeldt et al., 2011; Wilson et al., 2020; Wilson et al., 2021; Wilson et al., 2024). Among studies on wild birds (Benskin et al., 2009; Delgado-V and French, 2012), the impacts of human disturbances were not assessed. Some studies have documented how urbanization affects pathogen prevalence; however, they have primarily focused on terrestrial species and have rarely considered species’ diets (Bradley and Altizer, 2007; Murray et al., 2019). Predictions derived from such studies are unlikely to fully apply to seabirds, as they encounter different opportunities and constraints for pathogen transmission at sea and on land due to their unique ecology. Given the recent emergence of global wildlife pandemics, such as highly pathogenic avian influenza, linking human impacts to pathogen prevalence in seabirds can provide valuable insights for their conservation. We aim to fill the gap by addressing three questions: (1) whether the prevalence of pathogens in seabirds increases with human impact on marine and terrestrial environments, (2) if this association is influenced by an interaction with seabirds’ diets, and (3) whether pathogen associations with human disturbances vary by transmission route.

## 3 Materials and Methods

### 3.1 Literature review

We conducted a comprehensive literature search in Web of Science from 2000 to 2024 to identify English publications on seabirds and pathogens. The search terms were (seabird* OR gull* OR penguin* OR albatross* OR petrel* OR fulmar* OR shearwater* OR murre* OR tern* OR skua* OR puffin* OR frigatebird* OR auklet* OR guillemot* OR kittiwake* OR cormorant* OR shag* OR booby* OR tropicbird* OR auk* OR pelican* OR gannet*) AND (pathogen* OR disease* OR infection* OR parasite* OR virus* OR bacteria OR helminth* OR protozoa*). We restricted the search to keywords appearing in titles and abstracts to enhance relevance. We then screened titles and abstracts and excluded the ones that were irrelevant. To minimize bias, we excluded experimental studies and samples from captive birds, rehabilitation centers, and carcasses, as well as studies where the sampling effort relied on clinical signs and diagnoses were based on symptoms rather than laboratory tests. Additionally, we removed research on pooled samples, eggs, and studies that collected data from fecal samples from nests or the ground, rather than from specified individual birds, because the number of individuals was unknown in such cases. We excluded studies focused on vectors, rather than pathogens, and removed studies on shorebirds based on two criteria: species belonging to the families Scolopacidae (e.g., sandpipers), Charadriidae (e.g., plovers), Haematopodidae (e.g., oystercatchers), and Jacanidae (jacanas), and species that never occupy marine neritic or oceanic habitats at any stage of their life cycle (breeding or non-breeding) according to IUCN Red List (International Union for Conservation of Nature, 2025). We also removed datasets used in more than one study. In the final step, we excluded samples for which we could not obtain covariate values (primarily studies located far inland and thus lacking marine environment information).

From each study, we collected data on sample size, number of positive cases, geographic location (x and y coordinates if mentioned or coordinates extracted from Google Earth based on location name), pathogen, and detection method. If any of this data was missing in a study, we excluded that study as well. We categorized the methods for pathogen detection into two groups: 1) isolation, including methods such as DNA extraction, blood smears, physical examination, antigen detection, microscopy, culturing bacteria, and histological analysis, and 2) exposure, which includes different methods of detecting antibodies, such as Enzyme-Linked Immunosorbent Assay (ELISA) and Microscopic Agglutination Test (MAT).

For each sample, we recorded the counts of infected and uninfected individuals, the sampling year(s), geographic location, pathogen, and host species. Many studies contributed multiple samples because they reported several pathogens, species, locations, or, in a few cases, distinct sampling periods. So the number of samples in our dataset exceeds the number of studies (Figure S2 and Section S2). The literature search data are available in the Supplement Materials: Pathogen Prevalence in Seabirds Dataset.

### 3.2 Predictor variables

We used two spatial layers that quantify overall human disturbances that may affect pathogen prevalence: the Human Footprint Index for terrestrial environments (hereafter H.Land; Mu et al., 2022) and the Cumulative Human Impact on Marine Ecosystems dataset (hereafter H.Sea; Halpern et al., 2019). The H.Land layer combines eight layers of human pressure, including urban development, population density, nighttime lights, croplands, pasturelands, roads, railways, and navigable waterways. For 81 study locations that fell outside the H.Land layer (e.g., some remote small islands), we manually assigned scores by examining the corresponding areas using satellite imagery, following the methodology outlined in the original dataset (Mu et al., 2022; Venter et al., 2016). The H.Sea layer aggregates human impacts on global oceans from 19 anthropogenic stressors across four categories: land-based pollution (such as nutrient runoff); ocean-based activities (such as fishing); climate change; and invasive species. We created a buffer of 50 km around each data point and extracted the mean value for each covariate (H.Land and H.Sea) in each buffer. To simplify calculations and facilitate more explicit comparisons among predictors, we standardized these predictors using z-transformation (Beck et al., 2022).

According to the Pearson correlation coefficient, H.Land and H.Sea were not strongly colinear in our dataset (r = 0.40).

To understand the association between diet and its interaction with human impact, we extracted diet data from the EltonTraits 1.0 dataset (Wilman et al., 2014). We reclassified these diets as predominantly piscivore, predominantly invertivore, or generalist (Table 1). Species falling in the third category did not have a dominant diet except for two species that had unique diets compared to others: dolphin gull *Leucophaeus scoresbii* (dominantly scavenger) and brown skua *Stercorarius antarcticus* (mainly feeds on mammals and birds).

**TABLE 1.** Definition of each diet category extracted from EltonTraits 1.0 dataset (Wilman et al., 2014).

| Diet | Definition |
| --- | --- |
| Piscivore | $\geq 50\%$ of the diet is fish. |
| Invertivores | $\geq 50\%$ of the diet is invertebrates: aquatic invertebrates, shrimp, krill, squid, crustaceans, molluscs, cephalopods, polychaetes, gastropods, orthoptera, terrestrial invertebrates, ground insects, insect larvae, worms, orthopterans, and flying insects. |
| Generalists | None of the species-specific diets reach 50%, except for dolphin gull and brown skua. |

### 3.3 Extracting transmission route

To understand how the relationship between human disturbance and pathogen prevalence may be altered by its transmission route, we obtained data on pathogen transmission routes from vatious sources as we could not find one comprehensive database for pathogens transmission routes related to birds. We used the Global Mammal Parasite Database (GMPD), Centers for Disease Control and Prevention (CDC), and published articles related to pathogens not found in these two databases (Stephens et al., 2017; Centers for Disease Control and Prevention, 2025) (see Section S3). We classified these mechanisms into three categories: close and non-close contact (hereafter, contact-based), vector-borne, and intermediate host (Pedersen et al., 2005; Thrusfield, 2013). Close-contact pathogens spread through direct interactions (e.g., touching and biting) between individuals, while non-close contact pathogens spread indirectly through contaminated objects (e.g., surfaces) or environments (e.g., water, food sources, or soil) (Centers for Disease Control and Prevention, 2025). In this study, we combined these two groups because many viruses and bacteria in our dataset can be transmitted via both routes. Vector-borne pathogens rely on living carriers, typically arthropods such as mosquitoes, flies, and ticks (Centers for Disease Control and Prevention, 2025). The final category, intermediate host, contains pathogens that require a secondary host to complete part of their life cycle, such as growth or asexual reproduction, before passing to their definitive host. In our dataset, some pathogens belong to more than one category (Onstad et al., 2006).

### 3.4 Statistical analysis

Our analyses focused on identifying the factors that explain pathogen prevalence, defined as the number of infected individuals in each sample, bounded by the number of individuals tested for infection. Since many samples did not contain infected individuals, we used generalized linear mixed model (GLMM) with a zero-inflated binomial distribution (ZIB) using the R package glmmTMB (Britt, Rocque, and Zimmerman, 2018; Brooks et al., 2017; Hall, 2000). To account for the variability that our fixed effects cannot explain, we included two random effects. The first is the seabird species nested within genus, since the evolutionary history and genetic relatedness can influence a species’ susceptibility or resistance to a disease (Ilgunas et al., 2019). For example, birds in the order Passeriformes are more susceptible to avian malaria compared to other bird groups (Robles-Fernández, Santiago-Alarcon, and Lira-Noriega, 2022). Then, we incorporated a random effect for study ID to control for variation specific to each study.

We modeled pathogen prevalence (*Y_i_*) as a function of H.Land, H.Sea, and diet, and allowed excessive zeros to depend on the method of pathogen detection as

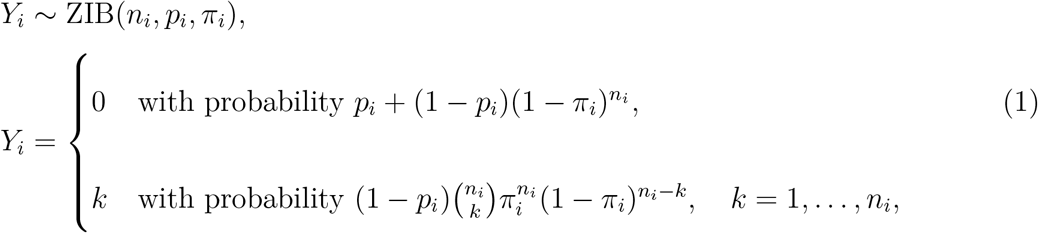

where *i* is a sampling unit defined with unique pathogen, seabird species, study, and location, *p_i_* is the probability of a zero because infection was not possible and/or not detectable (e.g., no exposure, unsuitable host, limits of the detection method; considered as the zero inflation), *n_i_* is the number of individuals in sample *i*, and *π_i_* is the probability of the individual being infected if infection is possible (i.e., probability of success) in a single trial (Hall, 2000).

The probability of individual infection is modelled as:

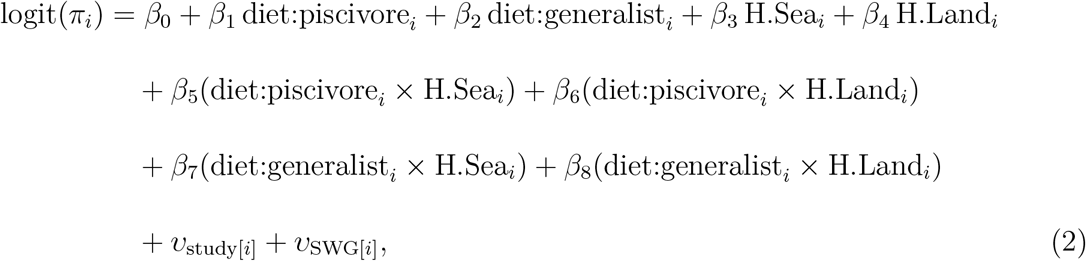

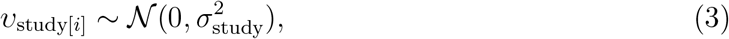

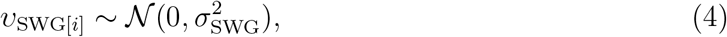

where *β_predictor_* is the coefficient for each predictor, *υ*_study[_*_i_*_]_ is random effect of each study for sample *i*, and *υ*_SBSwG[_*_i_*_]_ is random effect of seabird species within genus for sample *i*. While H.Sea*_i_* and H.Land*_i_* are continuous predictors, diet:piscivore*_i_* and diet:generalist*_i_* are dummy variables that identify whether the species in sample unit *i* fall in each of these diets. The probability of a structural zero that is not defined in the binomial distribution is modelled as

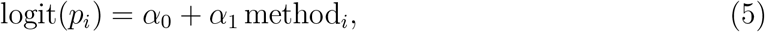

where *p_i_*is the probability that sample *i* is zero, *α*_0_ is the intercept, and *α*_1_ is the coefficient for the method of pathogen detection.

We conducted two sets of GLMMs to investigate differences between pathogen transmission routes. The first set included all available data, regardless of the pathogen transmission route (hereafter, the general model set). The second set comprised separate models focusing on each of the three transmission route categories (hereafter, contact-based, vector-borne, and intermediate host).

For model selection, we used the Bayesian Information Criterion (BIC) to identify the best-fitting models (Brewer, Butler, and Cooksley, 2016). To perform residual diagnostics, we applied simulation-based methods available in the DHARMa package (Hartig and Hartig, 2017). When the BIC-selected model showed any association with H.Sea or H.Land, we used parametric bootstrap likelihood-ratio tests to assess whether the model including H.Sea and H.Land significantly improved model fit relative to a nested model without these two covariates (for more detail, see section Parametric bootstrap likelihood ratio).

## 4 Results

Our literature-review dataset spans all oceans and includes samples from 5 orders, 35 seabird genera, and 65 speccies (Figure 1 and S5). The first three selected models by BIC (i.e., general, contact-based, and vector-borne models) retained both H.Sea and H.Land (Tables S1, S2, and S3), and parametric bootstrap likelihood-ratio tests showed adding human-related predictor can significantly improve the fit for the three models (general: Δ deviance = 78.2.9, *p*_value_ = 0.001; contact-based: Δ deviance = 101.2, *p*_value_ = 0.001; vector-borne: Δ deviance = 80.8, *p*_value_ = 0.001). The intermediate-host model showed no clear association with any predictors. DHARMa’s outlier test detected no extreme residuals, and the residual diagnostics indicated small but significant deviations for general, contact-based, and vector-borne models, and a poor fit for the intermediate-host model (Figure S3).

**FIGURE 1.**
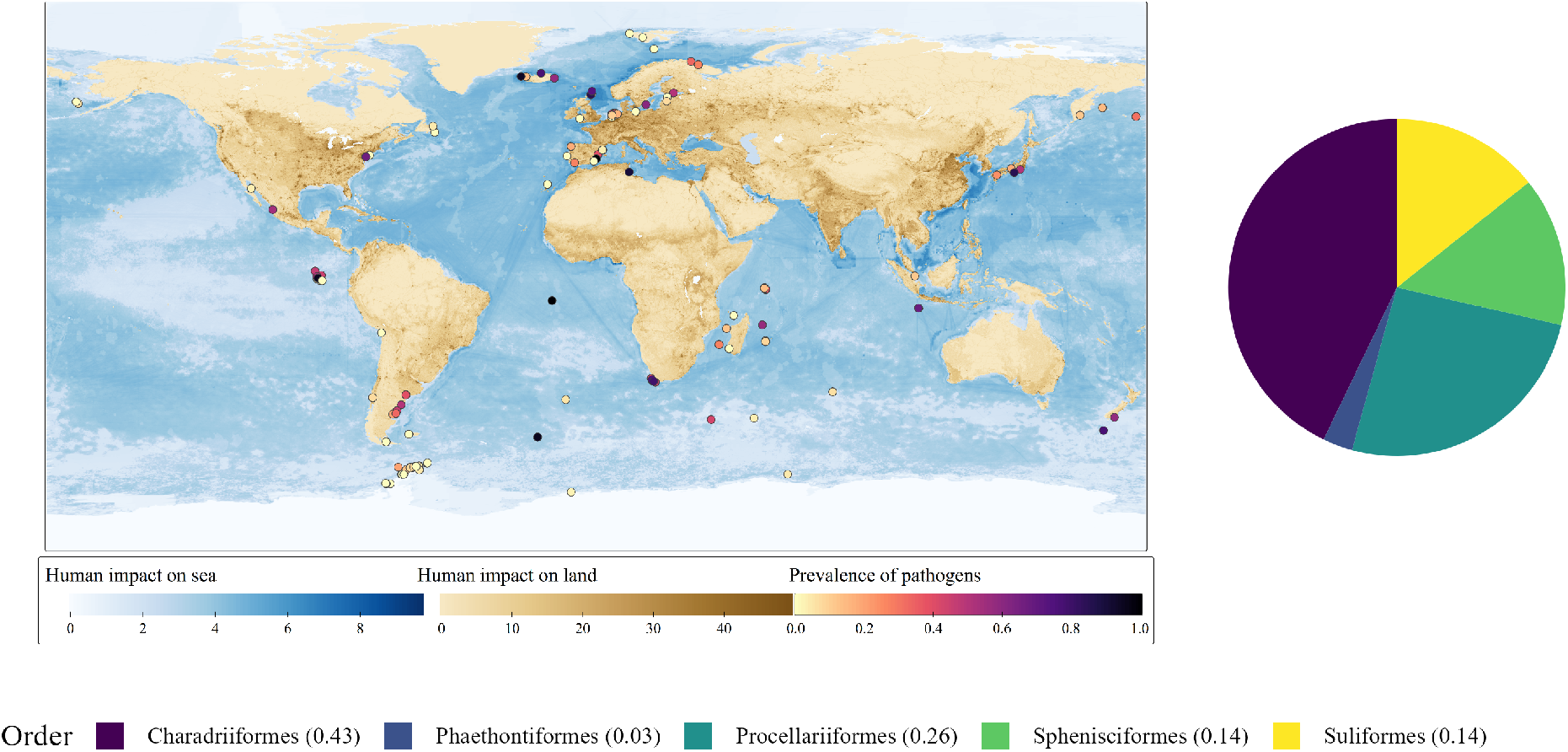
Global distribution of the samples and seabird orders. Each dot marks one independent dataset sample from published studies. The colour of each dot shows the pathogen prevalence reported in samples across a gradient of human impact on marine and terrestrial environments. The pie chart shows seabirds orders in the study. Our dataset contained 5 orders of seabirds with the most frequent being Charadriiformes.

Associations between pathogen prevalence and human impact varied with diet and transmission route (Figure 2 panel A; Table 2). The best model from the general set indicates that the prevalence in piscivores increased with both H.Sea and H.Land, and that the prevalence in generalists increased with H.Land. Invertivores, in contrast, showed a very small decline with increasing H.Land.

**FIGURE 2.**
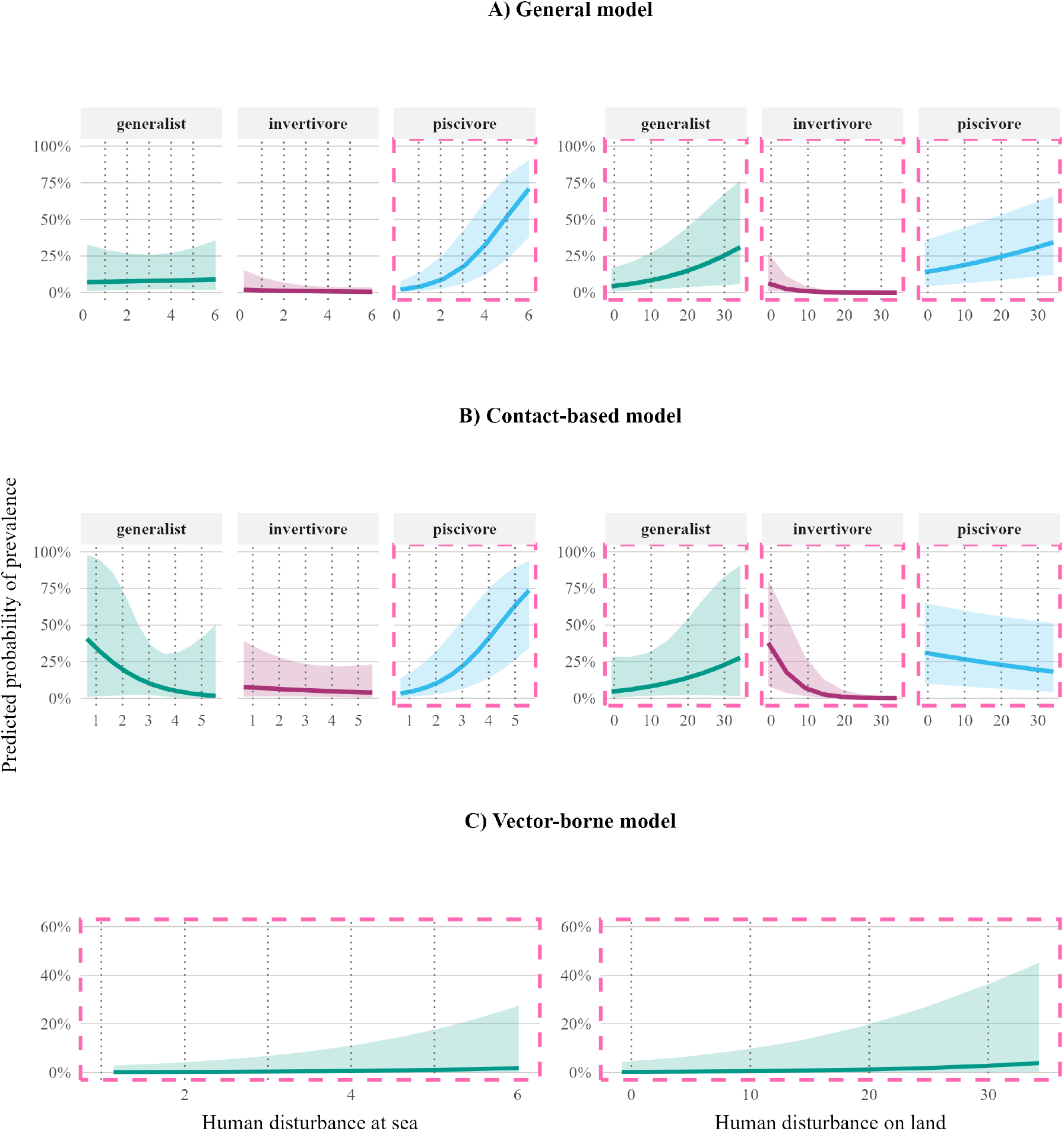
Pathogen prevalence as a function of H.Sea and H.Land and their interaction with 95% confidence intervals. The figures are A) general, B) contact-based, and C) vector-borne models. The main effects of H.Sea and H.Land give the slope for invertivores, and the interaction terms for generalists and piscivores show how their slopes differ from those of invertivores. Pink rectangles mark parameters whose 95% CIs do not include zero.

**TABLE 2.** Parameters in three BIC-selected models (General, Contact-based, Vector-borne). Estimates are on the logit scale. Here, FE and RE are fixed effects and random effects, respectively. The *p*-values in the first column are the result of the parametric bootstrap likelihood ratio test. The intercept corresponds to the reference diet (invertivore).

| Model<br>(p-value) | Term Variable | estimate | Standard<br>error | p-value |
| --- | --- | --- | --- | --- |
| General<br>(0.001) | FE intercept (invertivore) | $\beta_0 = -4.75$ | 0.84 | < 0.01 |
| | FE diet:piscivore | $\beta_1 = 3.2$ | 0.89 | < 0.01 |
| | FE diet:generalist | $\beta_2 = 2.21$ | 0.88 | 0.01 |
| | FE H.Sea (invertivore) | $\beta_3 = -0.23$ | 0.24 | 0.34 |
| | FE H.Land (invertivore) | $\beta_4 = -1.93$ | 0.49 | < 0.01 |
| | FE diet:piscivore × H.Sea | $\beta_5 = 1.02$ | 0.26 | < 0.01 |
| | FE diet:piscivore × H.Land | $\beta_6 = 2.26$ | 0.50 | < 0.01 |
| | FE diet:generalist × H.Sea | $\beta_7 = 0.26$ | 0.31 | 0.39 |
| | FE diet:generalist × H.Land | $\beta_8 = 2.59$ | 0.57 | < 0.01 |
| | RE Species within genus | $v_{\text{swg}} = 1.68$ | | |
| | RE Genus | $v_{\text{genus}} = 1.92$ | | |
| | RE Study | $v_{\text{study}} = 7.48$ | | |
| Contact-based<br>(0.001) | FE intercept (invertivore) | $\beta_0 = -2.55$ | 0.85 | < 0.01 |
| | FE diet:piscivore | $\beta_2 = 1.39$ | 0.99 | 0.16 |
| | FE diet:generalist | $\beta_1 = 0.22$ | 0.92 | 0.81 |
| | FE H.Sea (invertivore) | $\beta_3 = -0.15$ | 0.24 | 0.51 |
| | FE H.Land (invertivore) | $\beta_4 = -2.07$ | 0.49 | < 0.01 |
| | FE diet:piscivore × H.Sea | $\beta_5 = 1.04$ | 0.29 | < 0.01 |
| | FE diet:piscivore × H.Land | $\beta_6 = 1.88$ | 0.50 | < 0.01 |
| | FE diet:generalist × H.Sea | $\beta_7 = -0.54$ | 0.80 | 0.50 |
| | FE diet:generalist × H.Land | $\beta_8 = 2.68$ | 0.77 | < 0.01 |
| | RE Species within genus | $v_{\text{swg}} = 0.97$ | | |
| | RE Genus | $v_{\text{genus}} = 1.67$ | | |
| | RE Study | $v_{\text{study}} = 5.37$ | | |
| Vector-borne<br>(0.001) | FE intercept | $\beta_3 = -5.31$ | 1.51 | < 0.01 |
| | FE H.Sea | $\beta_3 = 0.51$ | 0.12 | < 0.01 |
| | FE H.Land | $\beta_4 = 0.82$ | 0.12 | < 0.01 |
| | RE Species within genus | $v_{\text{swg}} = 0.62$ | 0.38 | |
| | RE Genus | $v_{\text{genus}} = 3.38$ | | |
| | RE Study | $v_{\text{study}} = 2.44$ | | |

Best models from sets focused on specific transmission routes differed from the general results, indicating that transmission routes may alter the effect of human disturbances on pathogen prevalence (Figure 2 panels B nad C). As for the contact-based model, prevalence in piscivores increased with H.Sea, prevalence in generalists increased with H.Land, and prevalence of inventories decreased with H.Land. In contrast to the general model, pathogen prevalence in piscivores decreased with H.Land. However, the difference between the H.Land effect on piscivores and the baseline diet (invertivores) was positive. The prevalence of vector-borne pathogens was generally low, and showed small positive associations with both H.Sea and H.Land, regardless of diets (Figure 2). Other coefficients were not significant (their 95% confidence interval crossed zero; Figure S4). In the intermediate model, no covariate showed a significant association with prevalence. The intermediate dataset consisted of 85 samples from 4 studies. The small dataset may be why our models did not find any patterns.

To assess whether some seabird groups had higher overall pathogen prevalence than the average, we looked at the predicted odds ratio of pathogen prevalence for each genus. While most genera do not indicate clear deviation from the overall rate, two genera of seabirds (i.e., Pachyptila and Fregata) had odds ratios entirely above 1 in the selected vector-borne model (Figure S5).

## 5 Discussion

We found that pathogen prevalence in seabirds generally increased with human disturbance at sea (H.Sea) and on land (H.Land). Vector-borne pathogen prevalence rose with both H.Sea and H.Land across all diets, indicating a consistent positive association with disturbance. Diet and transmission route modified these patterns for other pathogen types. Piscivores showed particularly strong increases in prevalence with H.Sea, especially for contact-based pathogens (over 0.75 increase). Generalists had a higher prevalence with H.Land. In contrast, at high levels of H.Land, piscivores and invertivores showed lower prevalence of contact-based pathogens.

Human pressures can alter pathogen risks in wildlife through ecological and physiological mechanisms (e.g., Becker, Streicker, and Altizer, 2015; Werner and Nunn, 2020). Anthropogenic food leads to different disease outcomes in wildlife, often amplifying contact-based microparasites (e.g., many bacteria and viruses) by aggregating hosts at landfills, feeders, crops, or refuse (Becker, Streicker, and Altizer, 2015). Our results for generalist seabirds showed the same trend, with increases in pathogen prevalence with human disturbance on land. However, some studies showed contradictory results for macroparasites with complex life cycles. They suggest food provided by humans can shift away from infected intermediate hosts (Becker, Streicker, and Altizer, 2015; Werner and Nunn, 2020). Some studies did not show a consistent positive or negative effect of urbanisation on bird and mammal parasites. They argued that differences in prevalence can be a result of parasite types, and their transmission modes, and can increase with pollutants, chronic stress, and weak immune defences (Delgado-V and French, 2012; Bradley and Altizer, 2007; Jacobson et al., 2003; Martin et al., 2010; Rollins-Smith, 2017; Palmer, 2018). Our findings are consistent with this idea, as we also observed that the relationship between human disturbance and pathogen prevalence depends on the pathogen type and its transmission mode.

The increase in pathogen prevalence in piscivores in high human disturbance at sea (H.Sea) likely reflects their foraging ecology. Piscivores often are exposed to pathogens through their prey, as many fish serve as tolerant reservoirs for parasites (Novotny et al., 2004). Fish host many parasites to reach infective stages, so predation on fish efficiently passes on these parasites up the food chain (Marcogliese, 2002). Many marine helminths are generalists that move easily across species through food chains. At the same time, their long-lived larval stages persist in intermediate or paratenic hosts and keep infection pressure high (Marcogliese, 2002). Piscivorous seabirds often bio-accumulate contaminants as well, which can disrupt their endocrine and immune systems (Carravieri et al., 2020; Bracey et al., 2020; Sunderland et al., 2019; Whitney and Cristol, 2017), increasing the risk of infection. For instance, organochlorine loads correlate with immune suppression in Caspian terns (*Sterna caspia*) and toxic pollution and heavy metals such as polychlorinated biphenyls and mercury are correlated with higher prevalence of avian influenza in waterfowls (Grasman and Fox, 2001; Teitelbaum et al., 2022). The reduction in pathogen prevalence with increased disturbance on land (H.Land) association for contact-based pathogens for piscivores may result from their different behaviour in urban areas. Many seabirds are offshore species that breed on remote islets in dense colonies. However, some species are increasingly breeding in human-dominated landscapes, forming more dispersed colonies (e.g., on rooftops; Idle et al., 2021; Macfarlane, 1977; Clancy and Ward, 2020; Pichegru et al., 2024). Moreover, unlike generalists, there is limited evidence of strictly piscivorous seabirds aggregating on landfills and poultry facilities, where contact-based pathogens such as avian influenza can be transmitted. For example, a large-scale survey in the Netherlands (2006–2009) compared over 6000 birds across an area with varying gradients of urbanization, and found that avian influenza prevalence was higher in highly urbanized sites in comparison to low-urbanized sites (Verhagen et al., 2012). However, piscivorous seabirds (e.g., cormorants) inhabiting urban areas aggregate around anthropogenic fish-related resources, such as aquacultural sites and fishing piers (e.g., Burr et al., 2023; Thomas and Forys, 2022). Because our H.Land layer did not include aquaculture, it may underrepresent an important pathway of pathogen exposure for these species.

Generalist seabirds in cities frequently use landfills, agricultural areas, and other anthropogenic food sources (Ceia et al., 2023; Abell et al., 2008; Fuirst et al., 2018). Such locations can provide an opportunity for pathogen carriers to be in proximity with susceptible individuals (Miller et al., 2015; Fleischer et al., 2024). For example, ibises in Florida and gulls in the Iberian coast showed a higher prevalence of Salmonelosis and Campylobacter, respectively, in more developed areas originating from urban parks and human wastes (Hernandez et al., 2016; Seixas et al., 2025; Ramos et al., 2010). In addition, this behaviour can compromise their immune systems through poor-quality food (high in fat and low in protein) (Van Heugten, Coffey, and Spears, 1996; Coop and Kyriazakis, 2001; Ezenwa, 2004; Becker, Streicker, and Altizer, 2015). In an experimental study, mice with an anthropogenic diet showed lower ability to kill *Salmonella paratyphi* compared to the ones with a natural diet (Cummings et al., 2020).

Invertivores also showed a lower prevalence in highly disturbed lands. Invertivore species in our study are mostly offshore and have limited exposure to land-based pathogens, especially those transmitted through contact (BirdLife International, 2025). And similar to piscivorous seabirds, they often do not aggregate in highly contaminated sites on land. It is important to note that the accumulated H.Land we used did not account for sewage impact, which may lead to an underrepresentation of human disturbances in such locations. Invertivores foraging on estuarine mudflats contaminated by sewage can show a higher prevalence of pathogens. For example, Olrog’s gull feeding on benthic prey and crabs are exposed to pathogens originating from human disturbances (La Sala et al., 2013).

Our vector-borne category aggregates pathogens transmitted by a range of arthropods, such as ticks, fleas, louse flies, and mosquitoes. We found a consistent, though slight, positive association between human impact and diets. Climate change, warmer microclimates (e.g., in cities), and higher humidity can increase the number and survival of vectors such as mosquitoes and ticks (Descamps, 2013; Epstein, 2001; Campbell et al., 2002; Bradley and Altizer, 2007). For example, warmer winters in the Arctic are associated with higher prevalence and intensity of *Ixodes uriae* ticks in seabird colonies (Descamps et al., 2012). Recent work suggests an expansion or a change in the distribution of *I. uriae* towards higher latitude due to warmer winters in the Arctic (McCoy et al., 2023). Mosquitoes (Culex spp.) are also well adapted to man-made water bodies (e.g., treatment plants and sewers; Bradley, Gibbs, and Altizer, 2008). The abundance of mosquitoes can lead to increased transmission of vector-borne pathogens, such as West Nile virus in avian species (Bradley, Gibbs, and Altizer, 2008). Also, simplifying bird communities and favouring competent reservoir hosts may increase the prevalence of some vector-borne viruses (McCoy et al., 2023).

In our study, thin-billed prion (*Pachyptila belcheri*) and species within the genus Fregata (frigatebirds) showed higher odds of prevalence of vector-borne pathogens. A study showed that Haemoproteus *iwa* infections are particularly common in frigatebirds relative to other seabirds, and are transmitted by host-specialist hippoboscid flies *Olfersia spinifera* that are abundant on frigatebirds throughout their range in comparison to other species (Bastien et al., 2014). This result is in line with a review on seabirds (Quillfeldt et al., 2011). They argued that since frigatebirds breed in warm, nearshore tropical environments where vectors such as louse flies (Hippoboscidae) and biting midges (Ceratopogonidae) are abundant, they can experience a high exposure to such parasites as well (Quillfeldt et al., 2011). The same study also linked increased prevalence of another vector-borne pathogen, Plasmodium, to traits that prolong exposure at the nest, especially burrow nesting and longer fledging periods. *P. Belcheri* is a burrow-nester with a relatively long fledging period of about 50 days (Quillfeldt et al., 2007; Strange, 1980).

Many seabird species shift their diet between breeding and non-breeding seasons. Quantifying diet across the full annual cycle is difficult, especially during the non-breeding period, and most traditional diet studies are limited to the chick-rearing season (Einoder, 2009; Williams et al., 2007; Hedd et al., 2010). Therefore, our use of three coarse diet categories can introduce uncertainty. However, in the BIC model selection for general and contact-based pathogens, the best-supported models retained the interaction between diet and human disturbance, indicating that this covariate still captures significant variation in pathogen prevalence despite these limitations. Models without diet as a fixed effect had higher BIC values for these two models (Tables S1 to S4). The best vector-borne model result, however, did not include diet.

Our literature review identified important data gaps. We found a few studies on seabird pathogens with intermediate hosts that met our criteria. As a result, it was difficult to assess whether the lack of relationships between diet and human impacts was real or due to the small sample size. Pathogens requiring intermediate hosts can be influenced by human-modified environments. For instance, the prevalence of *T. gondii* in mammalian wildlife depends on their diet and habitat and increases with human population density (Wilson et al., 2021; Wilson et al., 2024). We also excluded many papers that lacked the necessary details about their data, such as fine-scale locations or counts of infected and uninfected individuals. We recommend future research to focus more on intermediate-host pathogens in seabirds and publish more detailed data, including diagnostic protocols, locations, sample sizes, and raw counts, to enable comparisons and meta-analyses. Overall, we saw that human impacts can increase the prevalence of pathogens in seabirds. One of the clearest patterns is related to piscivores and human disturance at sea. Our results showed that, through the low to high gradients of H.Sea, the prevalence of pathogens in seabirds can increase from nearly zero to around 70% and even higher in contact-based pathogens. Generalists exposed to high human disturbance on land have a higher rate of contact-transmitted pathogens. It is essential to reduce human pressure by lowering contaminants and enhancing waste treatment at sea, as well as improving waste management on land, with an understanding of how seabirds use highly disturbed landscape on land and sea. For example, landfill management and restricting access to seairds has shown an increase in their stress levels in the short term and controversial results due to losing a reliable food (e.g., Pereira et al., 2025; Zorrozua et al., 2020; Delgado et al., 2021). Identifying which combinations of diet and human disturbance drive the highest pathogen prevalence can be key to prioritizing monitoring and management efforts for seabird populations.

## 6 Acknowledgement

We are deeply grateful to the SȾÁUTW̱ (Tsawout) and W̱SIḴEM (Tseycum) for giving us access to X̱O, X̱DEȽ (Mandarte) Island. We thank Mark Hipfner, Peter Arcese, and Jade Baanstra for their assistance and collaboration throughout the study. This work was supported by Natural Sciences and Engineering Research Council of Canada (NSERC) Discovery Grant and Northern Research Supplement programs, Canada Research Chairs Tier II: Statistical Ecology, BC Knowledge Development fund, Canada Foundation for Innovation’s John R. Evans Leaders Fund, Cecil and Kathleen Morrow Scholarship, Werner and Hildegard Hesse Research Award in Ornithology award, President’s Academic Excellence Initiative PhD Awards, and the Four Year Fellowship (4YF).

## Supplementary Material

### S1 Materials and Methods

#### S1.1 Parametric bootstrap likelihood ratio

To test the significance of each alternative model, which includes the H.Sea and H.Land, we created a null model including only diet as a fixed effect, except for vector-borne model whose selected model, and therefore the null model, did no include diet. We used this fitted model to simulate 1,000 new datasets, and then for each simulated dataset, we refitted both null and alternative models fitted based on the BIC-selected model and computed the corresponding likelihood ratios. We calculated the p-value as (*k* + 1)/1001, where *k* is the number of simulated datasets in which the likelihood ratio was equal to or greater than that observed in the original dataset (Warmuth, Metzler, and Zamora-Gutierrez, 2023; North, Curtis, and Sham, 2002).

**FIGURE S1.**
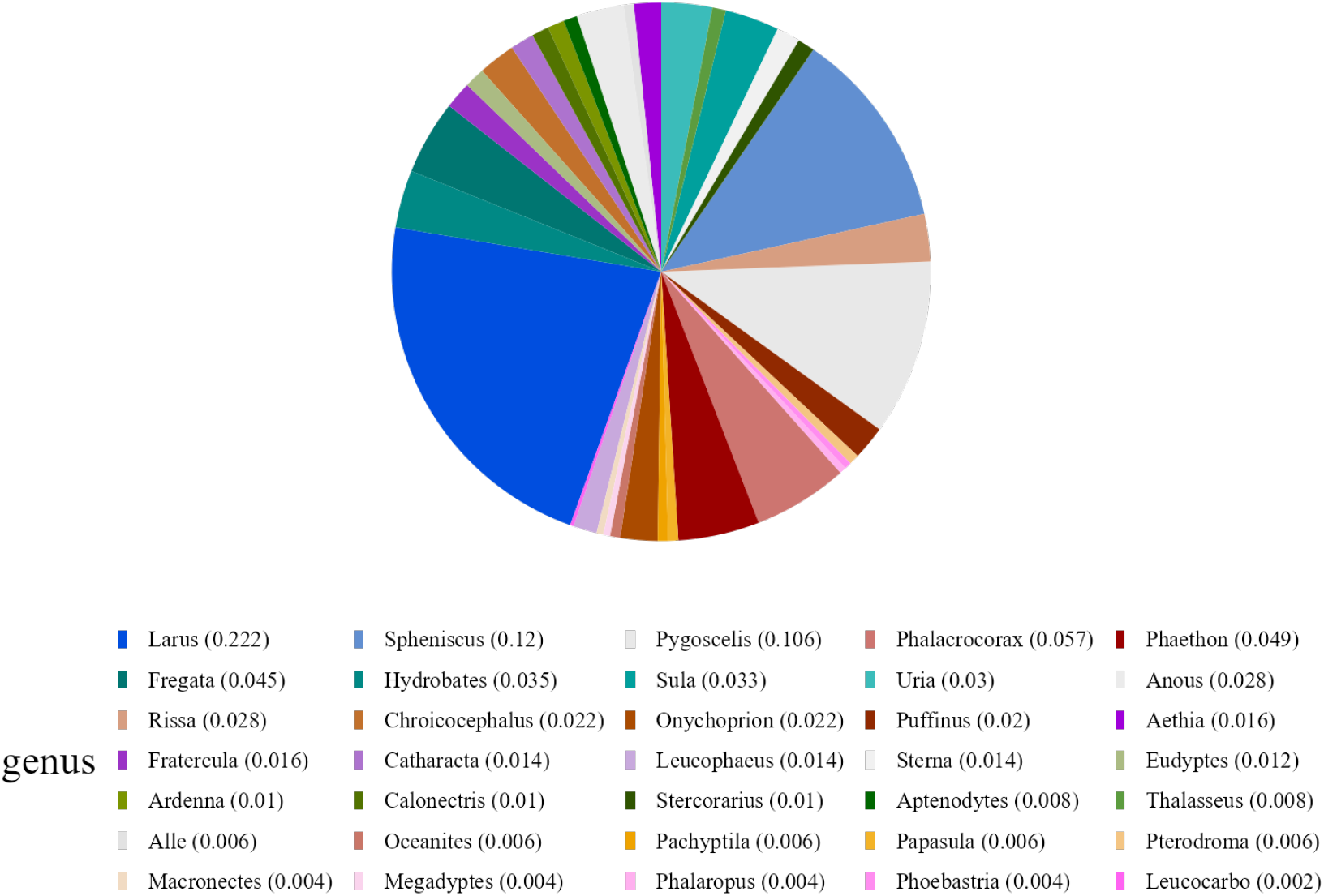
There are 35 genera of seabirds in our study. Gulls have the highest proportion (Larus 22%), followed by penguins (Spheniscus 11% and Pygoscelis 9%) and cormorants (Phalacrocorax 6%); all other genera each contributed 5% or less.

FIGURE S2. PRISMA flow diagram. We excluded studies lacking details on sample size, location, seabird species, or pathogen species. We also excluded study designs likely to bias our results, including experimental studies, samples from captive birds, rehabilitation centers, or carcasses, and studies where the number of individuals was not clear, and where sampling relied on clinical signs and diagnoses were based on symptoms rather than laboratory tests, and studies with sampling locations more than 50 km from the coastline. We collected 492 datasets from 54 studies, comprising 35 seabird genera and 65 species.

**TABLE S1.** The results of general BIC-selected model.

| Model # | Model (covariates) | BIC | ΔBIC |
| --- | --- | --- | --- |
| 1 | diet + H.Lnd + H.Mrn + diet:H.Lnd + diet:H.Mrn | 5254.6 | 0.00 |
| 2 | diet + H.Lnd + H.Mrn + diet:H.Lnd | 5260.3 | 5.75 |
| 3 | diet + H.Lnd + H.Mrn + diet:H.Lnd + diet:H.Mrn +<br>zi(mtC)* | 2560.6 | 6.05 |
| 4 | diet + H.Lnd + H.Mrn + diet:H.Mrn | 5265.0 | 10.44 |
| 5 | diet + H.Lnd + H.Mrn + diet:H.Mrn + zi(mtC) | 5271.1 | 16.51 |
| 6 | diet + H.Lnd + H.Mrn + diet:H.Lnd + zi(mtC) | 5271.9 | 17.34 |
| 7 | diet + H.Mrn + diet:H.Mrn | 5284.7 | 30.17 |
| 8 | diet + H.Mrn + diet:H.Mrn + zi(mtC) | 5289.8 | 35.23 |
| 9 | diet + H.Lnd + diet:H.Lnd + zi(mtC) | 5303.4 | 48.78 |
| 10 | H.Lnd | 5368.9 | 114.28 |
| 11 | (null) | 5368.6 | 115.01 |
| 12 | H.Marine | 5373.2 | 118.59 |
| 13 | diet + H.Lnd | 5373.9 | 119.36 |
| 14 | H.Lnd + H.Mrn | 5374.1 | 119.56 |
| 15 | diet | 5374.3 | 119.73 |
| 16 | H.Lnd + zi(mtC) | 5375.0 | 120.47 |
| 17 | diet + H.Marine + zi(mtC) | 5375.7 | 121.14 |
| 18 | diet + H.Lnd + H.Mrn | 5378.0 | 123.43 |
| 19 | H.Marine | 5379.3 | 124.68 |
| 20 | diet + H.Lnd + zi(mtC) | 5379.3 | 124.70 |
| 21 | H.Lnd + H.Marine + zi(mtC) | 5380.1 | 125.56 |
| 22 | diet + zi(mtC) | 5380.3 | 125.73 |
| 23 | H.Marine + zi(mtC) | 5380.5 | 125.89 |
| 24 | diet + H.Lnd + H.Mrn | 5384.1 | 129.56 |
| 25 | diet + H.Lnd + zi(mtC) | 5385.4 | 130.86 |
| 26 | diet + H.Lnd + diet:H.Lnd | - | - |

All models include conditional, dispersion, and zero-inflation intercepts.
\**zi(mtC)* is the method of pathogen detection in each sample in the zero-inflation.

**TABLE S2.** The results of contact-based BIC-selected model.

| Model # | Model (covariates) | BIC | ΔBIC |
| --- | --- | --- | --- |
| 1 | diet + H.Lnd + H.Mrn + diet:H.Lnd + diet:H.Mrn | 1592.5 | 0.00 |
| 2 | diet + H.Lnd + H.Mrn + diet:H.Lnd + diet:H.Mrn +<br>zi(mtC) | 1594.5 | 2.02 |
| 3 | diet + H.Mrn + diet:H.Mrn | 1599.1 | 6.56 |
| 4 | diet + H.Lnd + H.Mrn + diet:H.Mrn | 1599.2 | 6.65 |
| 5 | diet + H.Lnd + H.Mrn + diet:H.Lnd | 1601.0 | 8.46 |
| 6 | diet + H.Lnd + H.Mrn + diet:H.Lnd + zi(mtC) | 1601.1 | 8.58 |
| 7 | diet + H.Lnd + H.Mrn + diet:H.Mrn + zi(mtC) | 1601.2 | 8.67 |
| 8 | diet + H.Mrn + diet:H.Mrn + zi(mtC) | 1601.5 | 8.98 |
| 9 | diet + H.Lnd + diet:H.Lnd | 1607.6 | 15.07 |
| 10 | diet + H.Lnd + diet:H.Lnd + zi(mtC) | 1610.3 | 17.79 |
| 11 | H.Lnd | 1641.4 | 48.92 |
| 12 | H.Lnd + zi(mtC) | 1643.6 | 51.09 |
| 13 | H.Lnd + H.Mrn | 1645.6 | 53.07 |
| 14 | H.Lnd + H.Mrn + zi(mtC) | 1648.1 | 55.56 |
| 15 | H.Mrn + | 1651.1 | 58.60 |
| 16 | diet + H.Lnd | 1652.1 | 59.57 |
| 17 | (null) | 1652.5 | 16.01 |
| 18 | diet + H.Lnd + zi(mtC) | 1654.2 | 61.73 |
| 19 | H.Mrn | 1654.7 | 62.22 |
| 20 | zi(mtC) | 1655.4 | 62.93 |
| 21 | diet + H.Lnd + H.Mrn + zi(mtC) | 1656.2 | 63.69 |
| 22 | diet + H.Lnd + H.Mrn + zi(mtC) | 1658.7 | 66.19 |
| 23 | diet + H.Mrn | 1661.7 | 69.24 |
| 24 | diet | 1663.2 | 70.69 |
| 25 | diet + H.Mrn + zi(mtC) | 1665.4 | 72.87 |
| 26 | diet + zi(mtC) | - | - |

All models include conditional, dispersion, and zero-inflation intercepts.

**TABLE S3.**
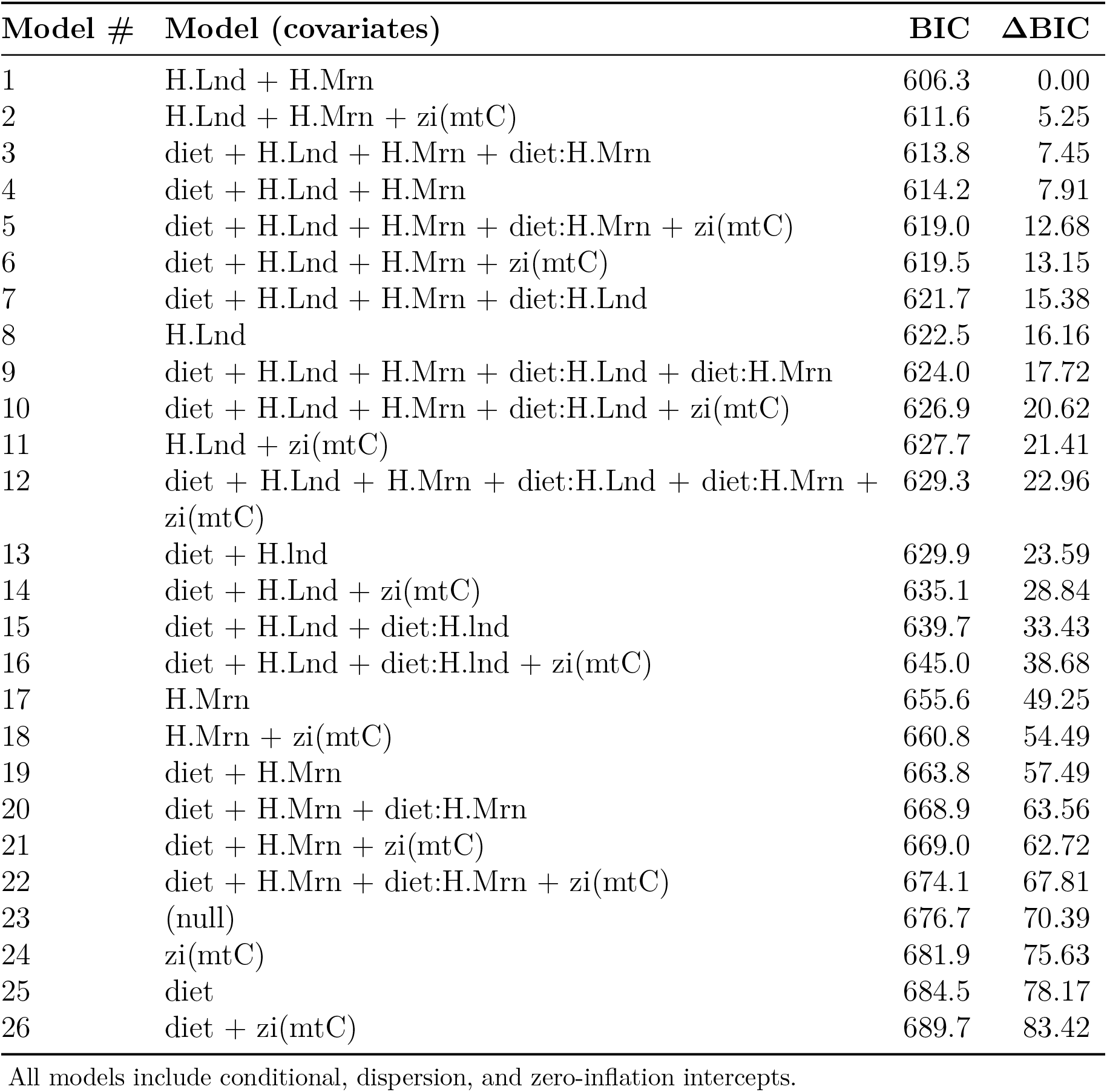
The results of vector-borne BIC-selected model.

**TABLE S4.**
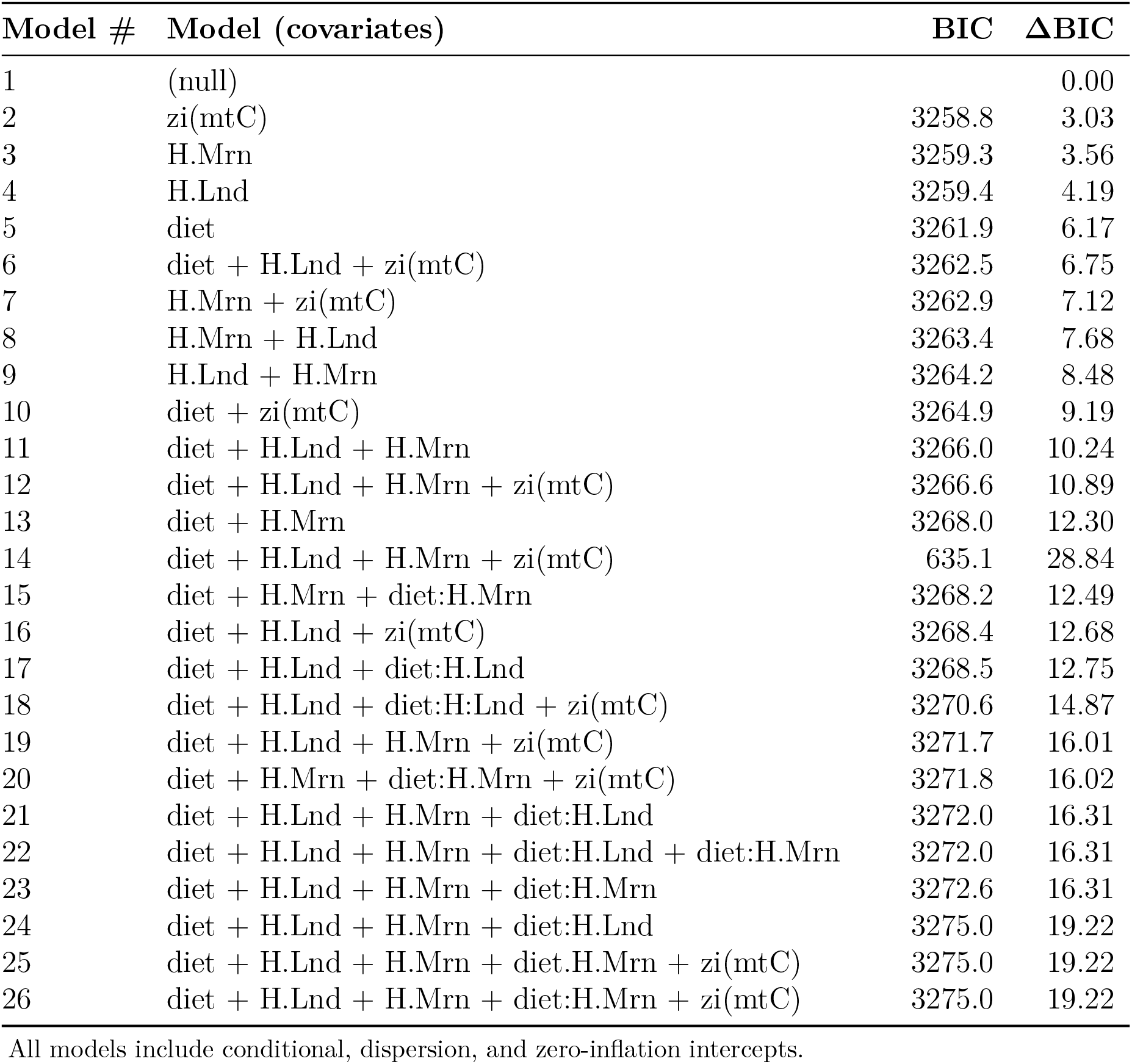
The results of Intermediate host BIC-selected model.

**FIGURE S3.**
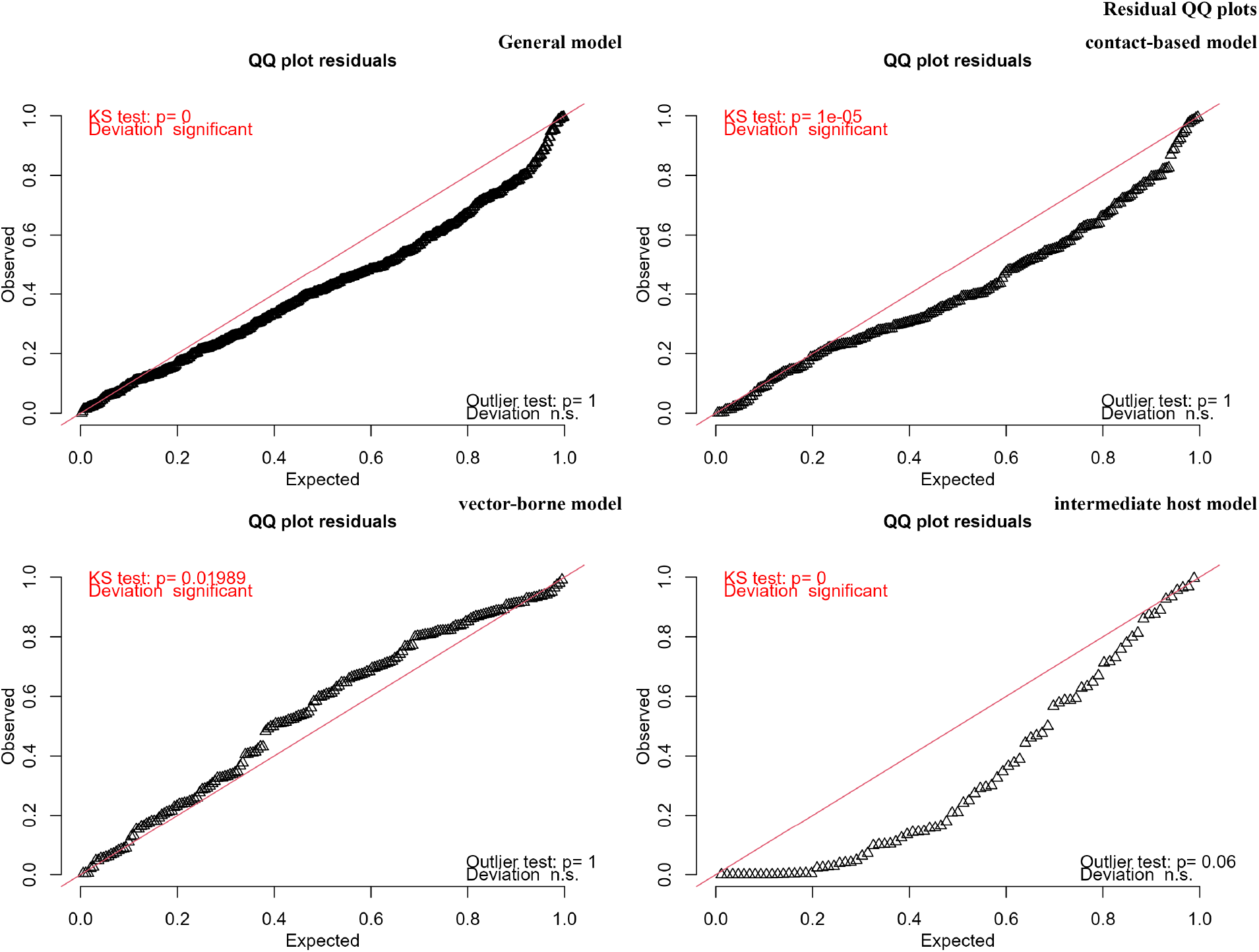
Residual diagnostics for prevalence models. DHARMa Q–Q plots of simulated residuals. Each panel reports the Kolmogorov–Smirnov tests. The general, contact-based, and vector-borne models show a small deviation, however the deviation for intermediate host model is large.

**FIGURE S4.**
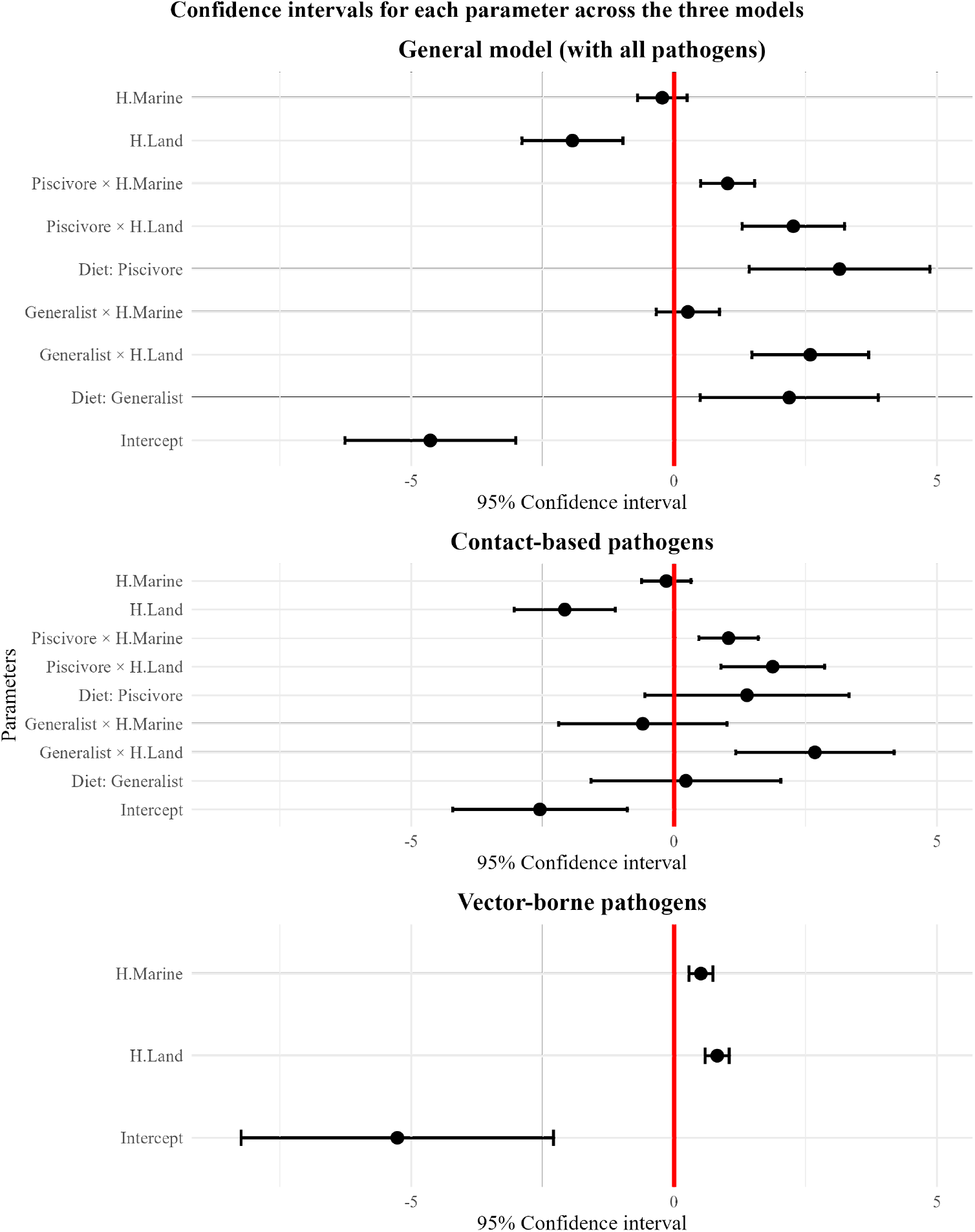
Predicted pathogen prevalence in seabirds by diet and human disturbance with 95% confidence interval.

**FIGURE S5.**
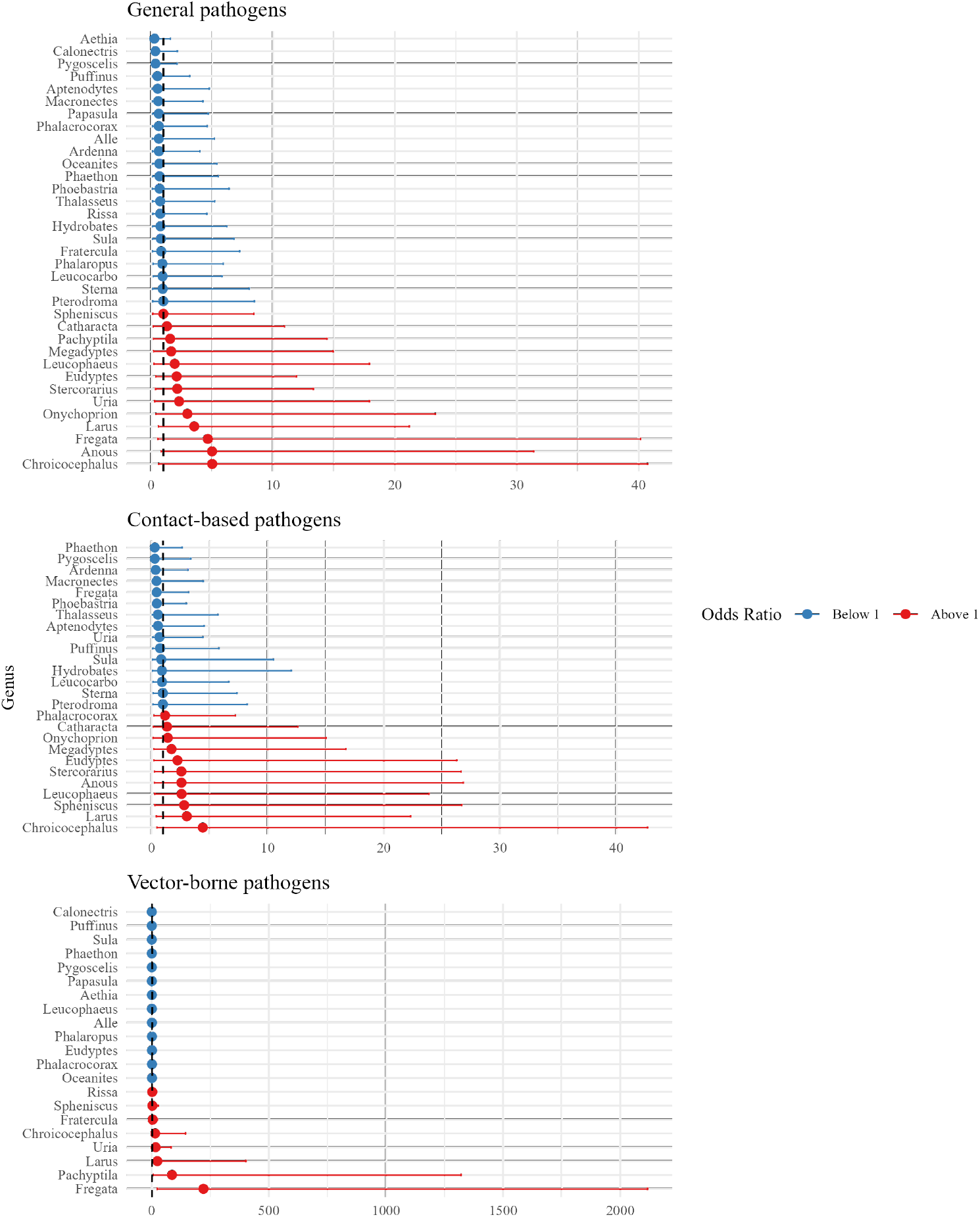
Odds ratio of seabird genera being infected with pathogens. Only two genera of Pachyptila and Fregata showed odds ratio over 1 with confidence interval not crossing 1 in vector-borne model. It means that these genera are more susceptible to vector-borne pathogens.

## S2 Meta analysis sources

## S3 Pathogen transmission route sources

## References

Abell, Robin, Michele L Thieme, Carmen Revenga, Mark Bryer, Maurice Kottelat, Nina Bogutskaya, Brian Coad, Nick Mandrak, Salvador Contreras Balderas, William Bussing, et al. 2008. “Freshwater ecoregions of the world: a new map of biogeographic units for freshwater biodiversity conservation.” Publisher: American Institute of Biological Sciences, BioScience 58 (5): 403–414.

Albery, Gregory F, Colin J Carlson, Lily E Cohen, Evan A Eskew, Rory Gibb, Sadie J Ryan, Amy R Sweeny, and Daniel J Becker. 2022. “Urban-adapted mammal species have more known pathogens.” Nature Ecology & Evolution 6 (6): 794–801.

Altizer, Sonia, Rebecca Bartel, and Barbara A Han. 2011. “Animal migration and infectious disease risk.” Science 331 (6015): 296–302.

Bartumeus, Frederic, Luca Giuggioli, Maite Louzao, Vincent Bretagnolle, Daniel Oro, and Simon A Levin. 2010. “Fishery discards impact on seabird movement patterns at regional scales.” Current Biology 20 (3): 215–222.

Bastien, Matthieu, Audrey Jaeger, Matthieu Le Corre, Pablo Tortosa, and Camille Lebarbenchon. 2014. “Haemoproteus iwa in great frigatebirds (*Fregata minor*) in the islands of the Western Indian Ocean.” PLoS One 9 (5): e97185.

Beck, Jared J, Daijiang Li, Sarah E Johnson, David Rogers, Kenneth M Cameron, Kenneth J Sytsma, Thomas J Givnish, and Donald M Waller. 2022. “Functional traits mediate individualistic species-environment distributions at broad spatial scales while fine-scale species associations remain unpredictable.” American Journal of Botany 109 (12): 1991–2005.

Becker, Daniel J, Daniel G Streicker, and Sonia Altizer. 2015. “Linking anthropogenic resources to wildlife–pathogen dynamics: a review and meta-analysis.” Ecology Letters 18 (5): 483–495.

Benskin, Clare McW H, Kenneth Wilson, Keith Jones, and Ian R Hartley. 2009. “Bacterial pathogens in wild birds: a review of the frequency and effects of infection.” Biological Reviews 84 (3): 349–373.

BirdLife International. 2025. IUCN Red List for birds. Accessed 2025-11-05. Accessed November 5, 2025. https://datazone.birdlife.org/.

Bracey, Annie M, Matthew A Etterson, Frederick C Strand, Sumner W Matteson, Gerald J Niemi, Francesca J Cuthbert, and Joel C Hoffman. 2020. “Foraging ecology differentiates life stages and mercury exposure in common terns (Sterna hirundo).” Integrated Environmental Assessment and Management 17 (2): 398–410.

Bradley, Catherine A, and Sonia Altizer. 2007. “Urbanization and the ecology of wildlife diseases.” Trends in ecology & evolution 22 (2): 95–102.

Bradley, Catherine A, Samantha EJ Gibbs, and Sonia Altizer. 2008. “Urban land use predicts West Nile virus exposure in songbirds.” Ecological Applications 18 (5): 1083–1092.

Bradley, Michael J, Susan J Kutz, Emily Jenkins, and Todd M O’Hara. 2005. “The potential impact of climate change on infectious diseases of Arctic fauna.” International Journal of Circumpolar Health 64 (5): 468–477.

Brearley, Grant, Jonathan Rhodes, Adrian Bradley, Greg Baxter, Leonie Seabrook, Daniel Lunney, Yan Liu, and Clive McAlpine. 2013. “Wildlife disease prevalence in human-modified landscapes.” Biological Reviews 88 (2): 427–442.

Brewer, Mark J, Adam Butler, and Susan L Cooksley. 2016. “The relative performance of AIC, AICC and BIC in the presence of unobserved heterogeneity.” Methods in Ecology and Evolution 7 (6): 679–692.

Britt, Chester L, Michael Rocque, and Gregory M Zimmerman. 2018. “The analysis of bounded count data in criminology.” Journal of Quantitative Criminology 34 (2): 591–607.

Brooks, Mollie E., Kasper Kristensen, Koen J. van Benthem, Arni Magnusson, Casper W. Berg, Anders Nielsen, Hans J. Skaug, Martin Maechler, and Benjamin M. Bolker. 2017. “glmmTMB balances speed and flexibility among packages for zero-inflated generalized linear mixed modeling.” The R Journal 9 (2): 378–400. https://journal.r-project.org/archive/2017/RJ-2017-066/.

Burr, Paul C, Brian S Dorr, Jimmy L Avery, Garrett M Street, and Bronson K Strickland. 2023. “Long term changes in aquaculture influence migration, regional abundance, and distribution of an avian species.” PloS One 18 (4): e0284265.

Campbell, Grant L, Anthony A Marfin, Robert S Lanciotti, and Duane J Gubler. 2002. “West nile virus.” The Lancet infectious diseases 2 (9): 519–529.

Camphuysen, CJ, SC Gear, and RW Furness. 2022. “Avian influenza leads to mass mortality of adult Great Skuas in Foula in summer 2022.” Scottish Birds 42:312–323.

Carlberg, Rachael A., Georgios Karris, Manish Verma, and Johannes Foufopoulos. 2022. “Food versus Disturbance: Contradictory Effects of Human Activities on an Opportunistic Seabird Breeding in an Oligotrophic Marine System.” Diversity 14 (6). issn: 14242818. 10.3390/d14060421.

Carr, Kelley S., and Michael S. Bronze. 2023. Vertical Transplacental Infections. https://www.ncbi.nlm.nih.gov/books/NBK604195/. Accessed: 2025-05-12.

Carravieri, Alice, Sarah J Burthe, Camille de La Vega, Yoshinari Yonehara, Francis Daunt, Mark A Newell, Rachel M Jeffreys, Alan J Lawlor, Alexander Hunt, Richard F Shore, et al. 2020. “Interactions between environmental contaminants and gastrointestinal parasites: novel insights from an integrative approach in a marine predator.” Environmental Science & Technology 54 (14): 8938–8948.

Ceia, Filipe R, Nathalie C Silva, Vitor H Paiva, Lurdes Morais, Ester A Serrão, and Jaime A Ramos. 2023. “Gulls as indicators of environmental changes in the North Atlantic: A long-term study on Berlenga Island, Western Portugal.” Diversity 15 (11): 1148.

Centers for Disease Control and Prevention. 2025. CDC website. https://www.cdc.gov. Accessed: 2025-09-04.

Cianchetti-Benedetti, Marco, G Dell’Omo, T Russo, C Catoni, and P Quillfeldt. 2018. “Interactions between commercial fishing vessels and a pelagic seabird in the southern Mediterranean Sea.” BMC Ecology 18:1–10.

Clancy, Cara, and Kim Ward. 2020. “Auto-rewilding in post-industrial cities: the case of inland cormorants in urban Britain.” Conservation and Society 18 (2): 126–136.

Cohen, Jeremy M, Erin L Sauer, Olivia Santiago, Samuel Spencer, and Jason R Rohr. 2020. “Divergent impacts of warming weather on wildlife disease risk across climates.” Science 370 (6519): eabb1702.

Coop, Robert L, and Ilias Kyriazakis. 2001. “Influence of host nutrition on the development and consequences of nematode parasitism in ruminants.” Trends in Parasitology 17 (7): 325–330.

Cortés-Hinojosa, Galaxia. 2024. “Marine bird of Neotropics, what we know, and we should know of diseases in a changing world.” In Ecology of Wildlife Diseases in the Neotropics, 121–141. Springer.

Cummings, Caroline R, Sonia M Hernandez, Maureen Murray, Taylor Ellison, Henry C Adams, Robert E Cooper, Shannon Curry, and Kristen J Navara. 2020. “Effects of an anthropogenic diet on indicators of physiological challenge and immunity of white ibis nestlings raised in captivity.” Ecology and Evolution 10 (15): 8416–8428.

Daszak, Peter, Andrew A Cunningham, and Alex D Hyatt. 2000. “Emerging infectious diseases of wildlife–threats to biodiversity and human health.” Science 287 (5452): 443–449.

Delgado, Sergio, Alfredo Herrero, A. Galarza, Asier Aldalur, Nere Zorrozua, and Juan Arizaga. 2021. “Demographic impact of landfill closure on a resident opportunistic gull.” Population Ecology 63 (3): 238–246. 10.1002/1438-390X.12083.

Delgado-V, Carlos A, and Kris French. 2012. “Parasite–bird interactions in urban areas: Current evidence and emerging questions.” Landscape and urban planning 105 (1-2): 5–14.

Descamps, Sébastien. 2013. “Winter temperature affects the prevalence of ticks in an Arctic seabird.” PLoS One 8 (6): e65374.

Descamps, Sebastien, Stephanie Jenouvrier, H Grant Gilchrist, and Mark R Forbes. 2012. “Avian cholera, a threat to the viability of an Arctic seabird colony?” PloS One 7 (2): e29659.

Dhama, K, M Mahendran, and S Tomar. 2008. “Pathogens transmitted by migratory birds: threat perceptions to poultry health and production.” International Journal of Poultry Science 7 (6): 516–525.

Dias, Maria P, Rob Martin, Elizabeth J Pearmain, Ian J Burfield, Cleo Small, Richard A Phillips, Oliver Yates, Ben Lascelles, Pablo Garcia Borboroglu, and John P Croxall. 2019. “Threats to seabirds: a global assessment.” Biological conservation 237:525–537.

Dietrich, Muriel, Elena Gómez-Díaz, and Karen D McCoy. 2011. “Worldwide distribution and diversity of seabird ticks: implications for the ecology and epidemiology of tick-borne pathogens.” Vector borne and zoonotic diseases 11 (5): 453–470.

Einoder, Luke D. 2009. “A review of the use of seabirds as indicators in fisheries and ecosystem management.” Fisheries Research 95 (1): 6–13.

Epstein, Paul R. 2001. “West Nile virus and the climate.” Journal of Urban Health 78 (2): 367–371.

Ezenwa, Vanessa O. 2004. “Interactions among host diet, nutritional status and gastrointestinal parasite infection in wild bovids.” International Journal for Parasitology 34 (4): 535–542.

Fackelmann, Gloria, Christopher K Pham, Yasmina Rodríguez, Mark L Mallory, Jennifer F Provencher, Julia E Baak, and Simone Sommer. 2023. “Current levels of microplastic pollution impact wild seabird gut microbiomes.” Nature Ecology & Evolution 7 (5): 698–706.

Fleischer, Ramona, Georg Joachim Eibner, Nina Isabell Schwensow, Fabian Pirzer, Sofia Paraskevopoulou, Gerd Mayer, Victor Max Corman, Christian Drosten, Kerstin Wilhelm, Alexander Christoph Heni, et al. 2024. “Immunogenetic-pathogen networks shrink in Tome’s spiny rat, a generalist rodent inhabiting disturbed landscapes.” Communications Biology 7 (1): 169.

Fuirst, Matthew, Richard R Veit, Megan Hahn, Nolwenn Dheilly, and Lesley H Thorne. 2018. “Effects of urbanization on the foraging ecology and microbiota of the generalist seabird Larus argentatus.” PLoS One 13 (12): e0209200.

Grasman, Keith A, and Glen A Fox. 2001. “Associations between altered immune function and organochlorine contamination in young Caspian terns (*Sterna caspia*) from Lake Huron, 1997–1999.” Ecotoxicology 10 (2): 101–114.

Grémillet, David, Aurore Ponchon, Pascal Provost, Amandine Gamble, Mouna Abed-Zahar, Alice Bernard, Nicolas Courbin, Grégoire Delavaud, Armel Deniau, Jérôme Fort, et al. 2023. “Strong breeding colony fidelity in northern gannets following high pathogenicity avian influenza virus (HPAIV) outbreak.” Biological Conservation 286:110269.

Grimaldi, Wray W, Phil J Seddon, Phil O’B Lyver, Shinichi Nakagawa, and Daniel M Tompkins. 2015. “Infectious diseases of Antarctic penguins: current status and future threats.” Polar Biology 38 (5): 591–606.

Gutiérrez, Jorge S., Eldar Rakhimberdiev, Theunis Piersma, and David W. Thieltges. 2017. “Migration and parasitism: habitat use, not migration distance, influences helminth species richness in Charadriiform birds.” _Eprint: https://onlinelibrary.wiley.com/doi/pdf/10.1111/jbi.12956, Journal of Biogeography 44 (5): 1137–1147. issn: 1365-2699, accessed May 3, 2024. 10.1111/jbi.12956.

Hall, Daniel B. 2000. “Zero-inflated Poisson and binomial regression with random effects: a case study.” Biometrics 56 (4): 1030–1039.

Halpern, Benjamin S., Melanie Frazier, Jamie Afflerbach, Julia S. Lowndes, Fiorenza Micheli, Casey O’Hara, Courtney Scarborough, and Kimberly A. Selkoe. 2019. “Recent pace of change in human impact on the world’s ocean.” Publisher: Nature Publishing Group, Scientific Reports 9, no. 1 (August 12, 2019): 11609. issn: 2045-2322, accessed March 5, 2024. 10.1038/s41598-019-47201-9.

Hartig, Florian, and Maintainer Florian Hartig. 2017. “Package ‘dharma’.” R package 531:532.

Hassell, James M, Michael Begon, Melissa J Ward, and Eric M Fèvre. 2017. “Urbanization and disease emergence: dynamics at the wildlife–livestock–human interface.” Trends in ecology & evolution 32 (1): 55–67.

Hedd, April, David A Fifield, Chantelle M Burke, William A Montevecchi, Laura McFarlane Tranquilla, Paul M Regular, Alejandro D Buren, and Gregory J Robertson. 2010. “Seasonal shift in the foraging niche of Atlantic puffins Fratercula arctica revealed by stable isotope (*δ*15N and *δ*13C) analyses.” Aquatic Biology 9 (1): 13–22.

Hernandez, Sonia M, Catharine N Welch, Valerie E Peters, Erin K Lipp, Shannon Curry, Michael J Yabsley, Susan Sanchez, Andrea Presotto, Peter Gerner-Smidt, Kelley B Hise, et al. 2016. “Urbanized white ibises (Eudocimus albus) as carriers of Salmonella enterica of significance to public health and wildlife.” PLoS One 11 (10): e0164402.

Holzman, David C. 2007. “Marine and Coastal Science: Where Space and Ocean Meet.” Publisher: National Institute of Environmental Health Sciences, Environmental Health Perspectives 115, no. 5 (May): A243. issn: 0091-6765, accessed April 25, 2023. 10.1289/EHP.115-A243A.

Hubálek, Zdeněk. 2021. “Pathogenic microorganisms associated with gulls and terns (Laridae).” Journal of Vertebrate Biology 70 (3): 21009–1.

Idle, Jessica L, Chad J Wilhite, Kristen C Harmon, Brooke Friswold, and Melissa R Price. 2021. “Wedge-tailed Shearwater (*Ardenna pacifica*) nesting success in human-dominated coastal environments.” PeerJ 9:e12096.

Ilgunas, Mikas, Vaidas Palinauskas, Elena Platonova, Tatjana Iezhova, and Gediminas Valkiunas. 2019. “The experimental study on susceptibility of common European songbirds to Plasmodium elongatum (lineage pGRW6), a widespread avian malaria parasite.” Malaria Journal 18 (1). issn: 14752875. 10.1186/s12936-019-2926-4.

International Union for Conservation of Nature. 2025. The IUCN Red List of Threatened Species. Accessed on 29 May 2025. https://www.iucnredlist.org.

Iverson, Samuel A, H Grant Gilchrist, Catherine Soos, Isabel I Buttler, N Jane Harms, and Mark R Forbes. 2016. “Injecting epidemiology into population viability analysis: avian cholera transmission dynamics at an arctic seabird colony.” Journal of Animal Ecology 85 (6): 1481–1490.

Jacobson, Kym C, Mary R Arkoosh, Anna N Kagley, Ethan R Clemons, Tracy K Collier, and Edmundo Casillas. 2003. “Cumulative effects of natural and anthropogenic stress on immune function and disease resistance in juvenile Chinook salmon.” Journal of Aquatic Animal Health 15 (1): 1–12.

Jara-Carrasco, S, M González, D González-Acuña, G Chiang, J Celis, W Espejo, P Mattatall, and R Barra. 2015. “Potential immunohaematological effects of persistent organic pollutants on chinstrap penguin.” Antarctic Science 27 (4): 373–381.

Jersey, Alix M de, Jennifer L Lavers, Alexander L Bond, Richard Wilson, Graeme R Zosky, and Jack Rivers-Auty. 2025. “Seabirds in crisis: Plastic ingestion induces proteomic signatures of multiorgan failure and neurodegeneration.” Science Advances 11 (11): eads0834.

Keesing, Felicia, Lisa K Belden, Peter Daszak, Andrew Dobson, C Drew Harvell, Robert D Holt, Peter Hudson, Anna Jolles, Kate E Jones, Charles E Mitchell, et al. 2010. “Impacts of biodiversity on the emergence and transmission of infectious diseases.” Nature 468 (7324): 647–652.

Khan, Junaid S., Jennifer F. Provencher, Mark R. Forbes, Mark L. Mallory, Camille Lebarbenchon, and Karen D. McCoy. 2019. “Parasites of seabirds: A survey of effects and ecological implications.” In Advances in Marine Biology, vol. 82. issn: 21625875. 10.1016/bs.amb.2019.02.001.

La Sala, Luciano F, Pablo F Petracci, Viviana Randazzo, and Mariano E Fernández-Miyakawa. 2013. “Enteric bacteria in Olrog’s gull (*Larus atlanticus*) and kelp gull (Larus dominicanus) from the Bahia Blanca Estuary, Argentina.” El hornero 28 (2): 59–64.

López-Calderón, Cosme, Víctor Martín-Vélez, Julio Blas, Ursula Höfle, Marta I Sánchez, Andrea Flack, Wolfgang Fiedler, Martin Wikelski, and Andy J Green. 2023. “White stork movements reveal the ecological connectivity between landfills and different habitats.” Movement Ecology 11 (1): 18.

Macfarlane, Anne E. 1977. “Roof-nesting by common terns.” Wilson Bulletin 89 (3): 22.

Marcogliese, DJ. 2002. “Food webs and the transmission of parasites to marine fish.” Parasitology 124 (7): 83–99.

Martin, Lynn B, William A Hopkins, Laura D Mydlarz, and Jason R Rohr. 2010. “The effects of anthropogenic global changes on immune functions and disease resistance.” Annals of the New York Academy of Sciences 1195 (1): 129–148.

Matos, Andressa Maria Rorato Nascimento de, Camila Domit, and Ana Paula Frederico Rodrigues Loureiro Bracarense. 2020. “Seabirds: studies with parasitofauna and potential indicator for environmental anthropogenic impacts.” Number: 4, Semina: Ciências Agrárias 41, no. 4 (May 13, 2020): 1439–1450. issn: 1679-0359, accessed February 27, 2024. 10.5433/1679-0359.2020v41n4p1439.

McCoy, Karen D, Muriel Dietrich, Audrey Jaeger, David A Wilkinson, Matthieu Bastien, Erwan Lagadec, Thierry Boulinier, Hervé Pascalis, Pablo Tortosa, Matthieu Le Corre, et al. 2016. “The role of seabirds of the Iles Eparses as reservoirs and disseminators of parasites and pathogens.” Acta Oecologica 72:98–109.

McCoy, Karen D, Céline Toty, Marlène Dupraz, Jérémy Tornos, Amandine Gamble, Romain Garnier, Sébastien Descamps, and Thierry Boulinier. 2023. “Climate change in the Arctic: Testing the poleward expansion of ticks and tick-borne diseases.” Global Change Biology 29 (7): 1729–1740.

McPhail, Gretchen M, Sydney M Collins, Tori V Burt, Noah G Careen, Parker B Doiron, Stephanie Avery-Gomm, Tatsiana Barychka, Matthew D English, Jolene A Giacinti, Megan EB Jones, et al. 2024. “Geographic, ecological, and temporal patterns of seabird mortality during the 2022 HPAI H5N1 outbreak on the island of Newfoundland.” Canadian Journal of Zoology 103:1–12.

Miller, Steven, Ulrike Zieger, Claudia Ganser, S Andrew Satterlee, Brittany Bankovich, Victor Amadi, Harry Hariharan, Diana Stone, and Samantha M Wisely. 2015. “Influence of land use and climate on Salmonella carrier status in the small Indian mongoose (*Herpestes auropunctatus*) in Grenada, West Indies.” Journal of Wildlife Diseases 51 (1): 60–68.

Morse, Stephen S, Jonna AK Mazet, Mark Woolhouse, Colin R Parrish, Dennis Carroll, William B Karesh, Carlos Zambrana-Torrelio, W Ian Lipkin, and Peter Daszak. 2012. “Prediction and prevention of the next pandemic zoonosis.” The Lancet 380 (9857): 1956–1965.

Mu, Haowei, Xuecao Li, Yanan Wen, Jianxi Huang, Peijun Du, Wei Su, Shuangxi Miao, and Mengqing Geng. 2022. “A global record of annual terrestrial Human Footprint dataset from 2000 to 2018.” Scientific Data 9 (1). issn: 20524463. 10.1038/s41597-022-01284-8.

Murray, Maureen H., Cecilia A. Sánchez, Daniel J. Becker, Kaylee A. Byers, Katherine E.L. Worsley-Tonks, and Meggan E. Craft. 2019. “City sicker? A meta-analysis of wildlife health and urbanization.” Frontiers in Ecology and the Environment 17 (10). issn: 15409309. 10.1002/fee.2126.

Newman, Scott H., Aleksei Chmura, Kathy Converse, A. Marm Kilpatrick, Nikkita Patel, Emily Lammers, and Peter Daszak. 2007. “Aquatic bird disease and mortality as an indicator of changing ecosystem health.” Marine Ecology Progress Series 352. issn: 01718630. 10.3354/meps07076.

North, Bernard V, David Curtis, and Pak C Sham. 2002. “A note on the calculation of empirical P values from Monte Carlo procedures.” The American Journal of Human Genetics 71 (2): 439–441.

Novotny, L, L Dvorska, A Lorencova, V Beran, I Pavlik, et al. 2004. “Fish: a potential source of bacterial pathogens for human beings.” Veterinární Medicína 49 (9): 343–358.

Onstad, DW, JR Fuxa, RA Humber, J Oestergaard, DI Shapiro-Ilan, VV Gouli, RS Anderson, TG Andreadis, and LA Lacey. 2006. “An abridged glossary of terms used in invertebrate pathology.” Society for Invertebrate Pathology.

Palmer, Caroline V. 2018. “Immunity and the coral crisis.” Communications Biology 1 (1): 91.

Pedersen, Amy B, Sonia Altizer, Mary Poss, Andrew A Cunningham, and Charles L Nunn. 2005. “Patterns of host specificity and transmission among parasites of wild primates.” International Journal for Parasitology 35 (6): 647–657.

Pereira, Jorge M, Jaime A Ramos, Adriana Domingues, Ana Almeida, Ana Marçalo, Catarina Cascão, Carlos Silva, Daniel Rey, Filipe R Ceia, Flávia Carvalho, et al. 2025. “Experimental anthropogenic food restrictions drive short-term foraging and immuno-haematological changes in sympatric breeding gulls.” Science of the Total Environment 1003:180672.

Pichegru, L, RB Sherley, T Malan, BJ Barham, K Ludynia, D Geldenhuys, K Amos, PJ Barham, E Drost, V Hahndiek, et al. 2024. “Decades of artificial nests towards African penguin conservation—Have they made a difference?” Ecological Solutions and Evidence 5 (4): e12388.

Plowright, Raina K, Patrick Foley, Hume E Field, Andy P Dobson, Janet E Foley, Peggy Eby, and Peter Daszak. 2011. “Urban habituation, ecological connectivity and epidemic dampening: the emergence of Hendra virus from flying foxes (Pteropus spp.)” Proceedings of the Royal Society B: Biological Sciences 278 (1725): 3703–3712.

Poulle, Marie-Lazarine, Matthieu Le Corre, Matthieu Bastien, Elsa Gedda, Chris Feare, Audrey Jaeger, Christine Larose, Nirmal Shah, Nina Voogt, Byron Göpper, et al. 2021. “Exposure of pelagic seabirds to *Toxoplasma gondii* in the Western Indian Ocean points to an open sea dispersal of this terrestrial parasite.” PLoS One 16 (8): e0255664.

Quillfeldt, Petra, Elena Arriero, Javier Martinez, Juan F Masello, and Santiago Merino. 2011. “Prevalence of blood parasites in seabirds-a review.” Frontiers in Zoology 8 (1): 26.

Quillfeldt, Petra, Ian J Strange, Gernot Segelbacher, and Juan F Masello. 2007. “Male and female contributions to provisioning rates of thin-billed prions, *Pachyptila belcheri*, in the South Atlantic.” Journal of Ornithology 148 (3): 367–372.

Ramos, Raül, Marta Cerdà-Cuéllar, Francisco Ramírez, Lluís Jover, and Xavier Ruiz. 2010. “Influence of refuse sites on the prevalence of Campylobacter spp. and Salmonella serovars in seagulls.” Applied and Environmental Microbiology 76 (9): 3052–3056.

Robles-Fernández, Ángel L., Diego Santiago-Alarcon, and Andrés Lira-Noriega. 2022. “Wildlife susceptibility to infectious diseases at global scales.” Proceedings of the National Academy of Sciences of the United States of America 119 (35). issn: 10916490. 10.1073/pnas.2122851119.

Rollins-Smith, Louise A. 2017. “Amphibian immunity–stress, disease, and climate change.” Developmental & Comparative Immunology 66:111–119.

Sadeghi, Sepideh, Mahnaz Nikaeen, Farzaneh Mohammadi, Amir Hossein Nafez, Sahar Gholipour, Zahra Shamsizadeh, and Mahdi Hadi. 2022. “Microbial characteristics of municipal solid waste compost: Occupational and public health risks from surface applied compost.” Waste Management 144. issn: 18792456. 10.1016/j.wasman.2022.03.012.

Sanz-Aguilar, Ana, Ana Payo-Payo, Andreu Rotger, Lena Yousfi, Sara Moutailler, Cecile Beck, Marine Dumarest, José Manuel Igual, Miguel Ángel Miranda, Mariana Viñas Torres, et al. 2020. “Infestation of small seabirds by *Ornithodoros maritimus* ticks: Effects on chick body condition, reproduction and associated infectious agents.” Ticks and Tick-borne diseases 11 (1): 101281.

Seixas, Julia Silva, Sonia M Hernandez, Katherine F Christie, William A Norfolk, R Scott Rozier, Elizabeth Kurimo-Beechuk, Lisa Hoopes, Michael J Yabsley, and Erin K Lipp. 2025. “Salmonella infections in white ibis nestlings in urban areas as a result of environmental contamination reveal a potential health risk to humans and animals.” Journal of the American Veterinary Medical Association 1 (aop): 1–8.

Simkin, Rohan D, Barbara A Han, Volker C Radeloff, Shannon LaDeau, Franz Schug, and Karen C Seto. 2025. “Zoonotic Host Richness in the Global Wildland–Urban Interface.” Global Change Biology 31 (2): e70039.

Stephens, Patrick R, Paula Pappalardo, Shan Huang, James E Byers, Maxwell J Farrell, Alyssa Gehman, Ria R Ghai, Sarah E Haas, Barbara Han, Andrew W Park, et al. 2017. Global mammal parasite database version 2.0.

Strange, Ian J. 1980. “The Thin-billed Prion, *Pachyptila belcheri*, at New Island, Falkand Islands [breeding sites; nest sites; behaviour; homing; predation, chick mortality].” Gerfaut 70.

Sunderland, Elsie M, Xindi C Hu, Clifton Dassuncao, Andrea K Tokranov, Charlotte C Wagner, and Joseph G Allen. 2019. “A review of the pathways of human exposure to poly-and perfluoroalkyl substances (PFASs) and present understanding of health effects.” Journal of Exposure Science & Environmental Epidemiology 29 (2): 131–147.

Teitelbaum, Claire S, Joshua T Ackerman, Mason A Hill, Jacqueline M Satter, Michael L Casazza, Susan EW De La Cruz, Walter M Boyce, Evan J Buck, John M Eadie, Mark P Herzog, et al. 2022. “Avian influenza antibody prevalence increases with mercury contamination in wild waterfowl.” Proceedings of the Royal Society B 289 (1982): 20221312.

Thomas, Fairl L, and Elizabeth A Forys. 2022. “The Role of Fishing Piers in Brown Pelican (*Pelecanus occidentalis*) Entanglement.” Animals 12 (18): 2352.

Thrusfield, Michael. 2013. Veterinary epidemiology. John Wiley & Sons.

Van Hemert, Caroline, John M Pearce, and Colleen M Handel. 2014. “Wildlife health in a rapidly changing North: focus on avian disease.” Frontiers in Ecology and the Environment 12 (10): 548–556.

Van Heugten, E, MT Coffey, and JW Spears. 1996. “Effects of immune challenge, dietary energy density, and source of energy on performance and immunity in weanling pigs.” Journal of Animal Science 74 (10): 2431–2440.

Venter, Oscar, Eric W. Sanderson, Ainhoa Magrach, James R. Allan, Jutta Beher, Kendall R. Jones, Hugh P. Possingham, et al. 2016. “Global terrestrial Human Footprint maps for 1993 and 2009.” Scientific Data 3. issn: 20524463. 10.1038/sdata.2016.67.

Verhagen, Josanne H, Vincent J Munster, Frank Majoor, Pascal Lexmond, Oanh Vuong, Job BG Stumpel, Guus F Rimmelzwaan, Albert DME Osterhaus, Martin Schutten, Roy Slaterus, et al. 2012. “Avian influenza a virus in wild birds in highly urbanized areas.” PLoS One 7 (6): e38256.

Warmuth, Vera M, Dirk Metzler, and Veronica Zamora-Gutierrez. 2023. “Human disturbance increases coronavirus prevalence in bats.” Science Advances 9 (13): eadd0688.

Werner, Courtney S, and Charles L Nunn. 2020. “Effect of urban habitat use on parasitism in mammals: a meta-analysis.” Proceedings of the Royal Society B 287 (1927): 20200397.

Whitney, Margaret C, and Daniel A Cristol. 2017. “Impacts of sublethal mercury exposure on birds: a detailed review.” Reviews of Environmental Contamination and Toxicology Volume 244, 113–163.

Williams, Cory T, C Loren Buck, Justine Sears, and Alexander S Kitaysky. 2007. “Effects of nutritional restriction on nitrogen and carbon stable isotopes in growing seabirds.” Oecologia 153 (1): 11–18.

Wilman, Hamish, Jonathan Belmaker, Jennifer Simpson, Carolina de la Rosa, Marcelo M. Rivadeneira, and Walter Jetz. 2014. “EltonTraits 1.0: Species-level foraging attributes of the world’s birds and mammals.” Ecology 95 (7). issn: 0012-9658. 10.1890/13-1917.1.

Wilson, Amy, David Lapen, Gregory Mitchell, Jennifer Provencher, and Scott Wilson. 2020. “Interaction of diet and habitat predicts *Toxoplasma gondii* infection rates in wild birds at a global scale.” Global Ecology and Biogeography 29 (7). issn: 14668238. 10.1111/geb.13096.

Wilson, Amy, David R Lapen, Jennifer F Provencher, and Scott Wilson. 2024. “The role of species ecology in predicting *Toxoplasma gondii* prevalence in wild and domesticated mammals globally.” PLoS Pathogens 20 (1): e1011908.

Wilson, Amy, Scott Wilson, Niloofar Alavi, and David R Lapen. 2021. “Human density is associated with the increased prevalence of a generalist zoonotic parasite in mammalian wildlife.” Proceedings of the Royal Society B 288 (1961): 20211724.

Zorrozua, Nere, Asier Aldalur, Alfredo Herrero, Diaz Beñat, Sergio Delgado, Carola Sanpera, Jover Lluís, and Juan Arizaga. 2020. “Breeding Yellow-legged Gulls increase consumption of terrestrial prey after landfill closure.” Ibis 162 (1): 50–62. 10.1111/ibi.12701.

## References

Abad, F. Xavier, Núria Busquets, Azucena Sanchez, Peter G. Ryan, Natàlia Majó, and Jacob Gonzalez-Solís. 2013. “Serological and virological surveys of the influenza A viruses in Antarctic and sub-Antarctic penguins.” Antarctic Science 25, no. 2 (March): 339–344. issn: 1365-2079. 10.1017/s0954102012001228.

Ariyama, Naomi, Rodrigo Tapia, Claudia Godoy, Belén Agüero, Valentina Valdés, Felipe Berrios, Pablo García Borboroglu, et al. 2021. “Avian orthoavulavirus 1 (Newcastle Disease virus) antibodies in five penguin species, Antarctic peninsula and Southern Patagonia.” Transboundary and Emerging Diseases 68, no. 6 (March): 3096–3102. issn: 1865-1682. 10.1111/tbed.14037.

Barriga, Gonzalo P., Dusan Boric-Bargetto, Marcelo Cortez San Martin, Víctor Neira, Harm van Bakel, Michele Thompsom, Rodrigo Tapia, et al. 2016. “Avian Influenza Virus H5 Strain with North American and Eurasian Lineage Genes in an Antarctic Penguin.” Emerging Infectious Diseases 22, no. 12 (December): 2221–2223. issn: 1080-6059. 10.3201/eid2212.161076.

Bennett-Laso, Benjamín, Bárbara Berazay, Gabriela Muñoz, Naomi Ariyama, Nikita Enciso, Christina Braun, Lucas Krüger, Miloš Barták, Marcelo González-Aravena, and Victor Neira. 2024. “Confirmation of highly pathogenic avian influenza H5N1 in skuas, Antarctica 2024.” Frontiers in Veterinary Science 11 (December). issn: 2297-1769. 10.3389/fvets.2024.1423404.

Bregnballe, Thomas, Christof Herrmann, Anja Globig, Anne Günther, Christoph Staubach, Joaquin Neumann Heise, Timm Harder, et al. 2024. “Outbreaks of highly pathogenic avian influenza (HPAI) epidemics in Baltic Great Cormorant *Phalacrocorax carbo* colonies in 2021 and 2022.” Bird Study 71, no. 4 (September): 353–366. issn: 1944-6705. 10.1080/00063657.2024.2399168.

Campioni, Letizia, Josué Martínez-de la Puente, Jordi Figuerola, José Pedro Granadeiro, Mónica C. Silva, and Paulo Catry. 2017. “Absence of haemosporidian parasite infections in the long-lived Cory’s shearwater: evidence from molecular analyses and review of the literature.” Parasitology Research 117, no. 1 (November): 323–329. issn: 1432-1955. 10.1007/s00436-017-5676-7.

Candela, Mónica G., Gonzalo G. Barberá, Angel Sallent, and Luis León. 2007. “Microbiological survey for selected bacterial pathogens in European storm petrel (*Hydrobates pelagicus*, Linnaeus 1758) from Grosa Island (Murcia, Southeastern Spain).” European Journal of Wildlife Research 54, no. 2 (November): 373–377. issn: 1439-0574. 10.1007/s10344-007-0147-6.

Contreras-Rodríguez, Araceli, Ma G. Aguilera-Arreola, Alma R. Osorio, María D. Martin, Rosa L. Guzmán, Enriqueta Velarde, and Enrico A. Ruiz. 2019. “Detection of Potential Human Pathogenic Bacteria Isolated From Feces of Two Colonial Seabirds Nesting on Isla Rasa, Gulf of California: Heermann’s Gull (*Larus heermanni*) and Elegant Tern *(Thalasseus elegans)*.” Tropical Conservation Science 12 (January). issn: 1940-0829. 10.1177/1940082919855673.

Espinaze, Marcela P. A., Cang Hui, Lauren Waller, Francois Dreyer, and Sonja Matthee. 2019. “Parasite diversity associated with African penguins *(Spheniscus demersus)* and the effect of host and environmental factors.” Parasitology 146, no. 6 (February): 791–804. issn: 1469-8161. 10.1017/s0031182018002159.

Fouchier, Ron A. M., Vincent Munster, Anders Wallensten, Theo M. Bestebroer, Sander Herfst, Derek Smith, Guus F. Rimmelzwaan, Björn Olsen, and Albert D. M. E. Osterhaus. 2005. “Characterization of a Novel Influenza A Virus Hemagglutinin Subtype (H16) Obtained from black-headed gulls.” Journal of Virology 79, no. 5 (March): 2814–2822. issn: 1098-5514. 10.1128/jvi.79.5.2814-2822.2005.

Fredes, Fernando, Cristian Madariaga, Eduardo Raffo, José Valencia, Marcela Herrera, Claudia Godoy, and Héctor Alcaíno. 2007. “Gastrointestinal parasite fauna of gentoo penguins *(Pygoscelis papua)* from the Península Munita, Bahía Paraíso, Antarctica.” Antarctic Science 19, no. 1 (February): 93–94. issn: 1365-2079. 10.1017/s0954102007000120.

Gallo Vaulet, Lucía, Ralph Eric Thijl Vanstreels, Luciana Gallo, Andrea Carolina Entrocassi, Laura Peker, Gabriela S. Blanco, Maria Virginia Rago, Marcelo Rodriguez Fermepin, and Marcela M. Uhart. 2022. “Chlamydiaceae-Like Bacterium in Wild Magellanic Penguins (Spheniscus magellanicus).” Diversity 14, no. 9 (September): 746. issn: 1424-2818. 10.3390/d14090746.

Gamble, Amandine, Romain Bazire, Karine Delord, Christophe Barbraud, Audrey Jaeger, Hubert Gantelet, Eric Thibault, et al. 2019. “Predator and scavenger movements among and within endangered seabird colonies: Opportunities for pathogen spread.” Edited by Sarah Knutie. Journal of Applied Ecology 57, no. 2 (December): 367–378. issn: 1365-2664. 10.1111/1365-2664.13531.

Gauthier-Clerc, Michel, Nicolas Eterradossi, Didier Toquin, Michèle Guittet, Grégoire Kuntz, and Yvon Le Maho. 2002. “Serological survey of the king penguin, *Aptenodytes patagonicus*, in Crozet Archipelago for antibodies to infectious bursal disease, influenza A and Newcastle disease viruses.” Polar Biology 25, no. 4 (April): 316–319. issn: 1432-2056. 10.1007/s00300-001-0346-7.

Guinn, Kayla, Alinde Fojtik, Nick Davis-Fields, Rebecca L. Poulson, Scott Krauss, Robert G. Webster, and David E. Stallknecht. 2016. “Antibodies to Influenza A Viruses in Gulls at Delaware Bay, USA.” Avian Diseases 60, no. 1s (May): 341–345. issn: 1938-4351. 10.1637/11103-042115-reg.

Hammouda, Abdessalem, Jessica Pearce-Duvet, Thierry Boulinier, and Slaheddine Selmi. 2014. “Egg sampling as a possible alternative to blood sampling when monitoring the exposure of yellow-legged gulls *(Larus michahellis)* to avian influenza viruses.” Avian Pathology 43, no. 6 (November): 547–551. issn: 1465-3338. 10.1080/03079457.2014.972340.

Hollmén, Tuula, J. Christian Franson, Douglas E. Docherty, Mikael Kilpi, Martti Hario, Lynn H. Creekmore, and Margaret R. Petersen. 2000. “Infectious bursal disease virus antibodies in eider ducks and herring gulls.” The Condor 102, no. 3 (August): 688–691. issn: 1938-5129. 10.1093/condor/102.3.688.

Jimenez-Uzcategui, Gustavo, Alberto Velez, Patricio Vega, Veronica Buendia, Verónica Montenegro-Benalcázar, Christian Sevilla, and Marilyn Cruz. 2024. “No evidence for high pathogenicity avian influenza in Waved Albatross (*Pvhoebastria irrorata*).” Marine Ornithology 52 (2): 349–353.

Kay, Emily, Melanie J. Young, Chris Muller, Laryssa Howe, Wendi Roe, and Brett D. Gartrell. 2022. “Prevalence and pathogen load of Eimeria in wild yellow-eyed penguins *(Megadyptes antipodes)* and the morphologic characterization of a novel Eimeria species.” Journal of Wildlife Diseases 58, no. 4 (November). issn: 0090-3558. 10.7589/jwd-d-21-00146.

Kim, Han-Kyu, Chang-Yong Choi, Min-Su Jeong, Hwa-Yeon Kang, and Woo-Shin Lee. 2016. “Regurgitation of the koilin layer in chinstrap penguins (*Pygoscelis antarcticus*) and its association with gastric parasites.” Polar Research 35, no. 1 (January): 25966. issn: 1751-8369. 10.3402/polar.v35.25966.

Kleinertz, S., S. Christmann, L. M. R. Silva, J. Hirzmann, C. Hermosilla, and A. Taubert. 2014. “Gastrointestinal parasite fauna of Emperor Penguins *(Aptenodytes forsteri)* at the Atka Bay, Antarctica.” Parasitology Research 113, no. 11 (August): 4133–4139. issn: 1432-1955. 10.1007/s00436-014-4085-4.

Krams, Indrikis, Valērija Suraka, Kalev Rattiste, Mikus Āboliņš-Ābols, Tatjana Krama, Markus J. Rantala, Pranas Mierauskas, Dina Cīrule, and Lauri Saks. 2012. “Comparative analysis reveals a possible immunity-related absence of blood parasites in Common Gulls *(Larus canus)* and Black-headed Gulls *(Chroicocephalus ridibundus)*.” Journal of Ornithology 153, no. 4 (May): 1245–1252. issn: 2193-7206. 10.1007/s10336-012-0859-6.

Kuklin, V. V., and M. M. Kuklina. 2021. “Age Dynamics of Helminth Fauna of the Herring Gull (*Larus argentatus*) in Kola Bay, Barents Sea.” Biology Bulletin 48, no. 8 (December): 1160–1169. issn: 1608-3059. 10.1134/s1062359021080173.

Lamb, Juliet, Jeremy Tornos, Romain Dedet, Hubert Gantelet, Nicolas Keck, Juliette Baron, Marine Bely, et al. 2022. “Hanging out at the club: Breeding status and territoriality affect individual space use, multi-species overlap and pathogen transmission risk at a seabird colony.” Functional Ecology 37, no. 3 (December): 576–590. issn: 1365-2435. 10.1111/1365-2435.14240.

Lebarbenchon, Camille, Audrey Jaeger, Matthieu Bastien, Matthieu Le Corre, Koussay Dellagi, and Hervé Pascalis. 2013. “Absence of Coronaviruses, Paramyxoviruses, and Influenza A Viruses in Seabirds in the Southwestern Indian Ocean.” Journal of Wildlife Diseases 49, no. 4 (October): 1056–1059. issn: 0090-3558. 10.7589/2012-09-227.

Lebarbenchon, Camille, Audrey Jaeger, Chris Feare, Matthieu Bastien, Muriel Dietrich, Christine Larose, Erwan Lagadec, et al. 2015. “Influenza A virus on oceanic islands: Host and viral diversity in seabirds in the western Indian Ocean.” Edited by Daniel R. Perez. PLoS Pathogens 11, no. 5 (May): e1004925. issn: 1553-7374. 10.1371/journal.ppat.1004925.

Levin, Iris I., Diana C. Outlaw, F. Hernán Vargas, and Patricia G. Parker. 2009. “Plasmodium blood parasite found in endangered Galapagos penguins *(Spheniscus mendiculus)*.” Biological Conservation 142, no. 12 (December): 3191–3195. issn: 0006-3207. 10.1016/j.biocon.2009.06.017.

Levin, Iris I., and Patricia G. Parker. 2012. “Prevalence of Hemoproteus iwa in Galapagos Great Frigatebirds *(Fregata minor)* and Their Obligate Fly Ectoparasite *(Olfersia spinifera)*.” Journal of Parasitology 98, no. 5 (October): 924–929. issn: 1937-2345. 10.1645/ge-3027.1.

Lobato, Elisa, Jessica Pearce-Duvet, Vincent Staszewski, Elena Gomez-Diaz, Jacob González-Solís, Alexander Kitaysky, Karen D McCoy, and Thierry Boulinier. 2011. “Seabirds and the circulation of Lyme borreliosis bacteria in the North Pacific.” Vector-Borne and Zoonotic Diseases 11 (12): 1521–1527.

Lobato, Elisa, Jessica Pearce-Duvet, Vincent Staszewski, Elena Gómez-Díaz, Jacob González-Solís, Alexander Kitaysky, Karen D. McCoy, and Thierry Boulinier. 2011. “Seabirds and the Circulation of Lyme Borreliosis Bacteria in the North Pacific.” Vector-Borne and Zoonotic Diseases 11, no. 12 (December): 1521–1527. issn: 1557-7759. 10.1089/vbz.2010.0267.

Martín-Vélez, Víctor, Joan Navarro, Jordi Figuerola, Raül Aymí, Sara Sabaté, Raquel Planell, Jordi Vila, and Tomás Montalvo. 2024. “A spatial analysis of urban gulls contribution to the potential spread of zoonotic and antibiotic-resistant bacteria.” Science of The Total Environment 912 (February): 168762. issn: 0048-9697. 10.1016/j.scitotenv.2023.168762.

Mendes, Luisa, Theunis Piersma, Miguel Lecoq, Bernard Spaans, and Robert E. Ricklefs. 2005. “Disease-limited distributions? Contrasts in the prevalence of avian malaria in shorebird species using marine and freshwater habitats.” Oikos 109, no. 2 (March): 396–404. issn: 1600-0706. 10.1111/j.0030-1299.2005.13509.x.

Merino, Santiago, Janos Hennicke, Javier Martínez, Katrin Ludynia, Roxana Torres, Thierry M. Work, Stedson Stroud, Juan F. Masello, and Petra Quillfeldt. 2012. “Infection by Haemoproteus Parasites in Four Species of Frigatebirds and the Description of a New Species of Haemoproteus (*Haemosporida: Haemoproteidae*).” Journal of Parasitology 98, no. 2 (April): 388–397. issn: 1937-2345. 10.1645/ge-2415.1.

Navarro, Joan, David Grémillet, Isabel Afán, Francisco Miranda, Willem Bouten, Manuela G. Forero, and Jordi Figuerola. 2019. “Pathogen transmission risk by opportunistic gulls moving across human landscapes.” Scientific Reports 9, no. 1 (July). issn: 2045-2322. 10.1038/s41598-019-46326-1.

Odoi, Justice Opare, Michiyo Sugiyama, Yuko Kitamura, Akiko Sudo, Tsutomu Omatsu, and Tetsuo Asai. 2021. “Prevalence of antimicrobial resistance in bacteria isolated from Great Cormorants *(Phalacrocorax carbo hanedae)* in Japan.” Journal of Veterinary Medical Science 83 (8): 1191–1195.

Pedro, Rodrigues, Navarrete Claudio, Campos Elena, and Verdugo Claudio. 2018. “Low occurrence of hemosporidian parasites in the Neotropic cormorant (*Phalacrocorax brasilianus*) in Chile.” Parasitology Research 118, no. 1 (November): 325–333. issn: 1432-1955. 10.1007/s00436-018-6146-6.

Poulle, Marie-Lazarine, Matthieu Le Corre, Matthieu Bastien, Elsa Gedda, Chris Feare, Audrey Jaeger, Christine Larose, et al. 2021. “Exposure of pelagic seabirds to *Toxoplasma gondii* in the Western Indian Ocean points to an open sea dispersal of this terrestrial parasite.” Edited by Vitor Hugo Rodrigues Paiva. PLoS One 16, no. 8 (August): e0255664. issn: 1932-6203. 10.1371/journal.pone.0255664.

Quillfeldt, Petra, Javier Martínez, Janos Hennicke, Katrin Ludynia, Anja Gladbach, Juan F. Masello, Samuel Riou, and Santiago Merino. 2010. “Hemosporidian blood parasites in seabirds—a comparative genetic study of species from Antarctic to tropical habitats.” Naturwissenschaften 97, no. 9 (July): 809–817. issn: 1432-1904. 10.1007/s00114-010-0698-3.

Rivera, William A., Jusuf S. Husic, Corey E. Gaylets, Roy H. Mosher, Brian G. Palestis, and Adam J. Houlihan. 2012. “Carriage of bacteria and protozoa in the intestinal tract of common tern chicks.” Waterbirds 35, no. 3 (September): 490–494. issn: 1938-5390. 10.1675/063.035.0314.

Rohrer, Sage D., Gustavo Jiménez-Uzcátegui, Patricia G. Parker, and Lon M. Chubiz. 2023. “Composition and function of the Galapagos penguin gut microbiome vary with age, location, and a putative bacterial pathogen.” Scientific Reports 13, no. 1 (April). issn: 2045-2322. 10.1038/s41598-023-31826-y.

Sanmartín, M.L., J.A. Cordeiro, M.F. Álvarez, and J. Leiro. 2005. “Helminth fauna of the yellow-legged gullLarus cachinnansin Galicia, north-west Spain.” Journal of Helminthology 79, no. 4 (December): 361–371. issn: 1475-2697. 10.1079/joh2005309.

Souza Petersen Elisa, de, de Araujo Jansen, Krüger Lucas, Seixas Marina M., Ometto Tatiana, Thomazelli Luciano M., Walker David, Durigon Edison Luiz, and Petry Maria Virginia. 2017. “First detection of avian influenza virus (H4N7) in giant petrel monitored by geolocators in the Antarctic region.” Marine Biology 164 (4). issn: 1432-1793. 10.1007/s00227-017-3086-0.

Staszewski, V, K.D McCoy, and T Boulinier. 2008. “Variable exposure and immunological response to Lyme disease Borrelia among North Atlantic seabird species.” Proceedings of the Royal Society B: Biological Sciences 275, no. 1647 (June): 2101–2109. issn: 1471-2954. 10.1098/rspb.2008.0515.

Ushine, Nana, Makoto Ozawa, Shouta MM Nakayama, Mayumi Ishizuka, Takuya Kato, and Shin-ichi Hayama. 2023. “Evaluation of the effect of Pb pollution on avian influenza virus-specific antibody production in black-headed gulls (*Chroicocephalus ridibundus*).” Animals 13 (14): 2338.

Vanstreels, Ralph Eric Thijl, Flavia R. Miranda, Valeria Ruoppolo, Ana Olívia de Almeida Reis, Erli Schneider Costa, Adriana Rodrigues de Lira Pessôa, João Paulo Machado Torres, et al. 2013. “Investigation of blood parasites of pygoscelid penguins at the King George and Elephant Islands, South Shetlands Archipelago, Antarctica.” Polar Biology 37, no. 1 (August): 135–139. issn: 1432-2056. 10.1007/s00300-013-1401-x.

Vanstreels, Ralph Eric Thijl, Marcela Uhart, Virginia Rago, Renata Hurtado, Sabrina Epiphanio, and Jose Luiz Catao-Dias. 2017. “Do blood parasites infect Magellanic penguins *(Spheniscus magellanicus)* in the wild? Prospective investigation and climatogeographic considerations.” Parasitology 144 (5): 698–705.

Verdugo, Claudio, Adrián Pinto, Naomi Ariyama, Manuel Moroni, and Carlos Hernandez. 2019. “Molecular identification of avian viruses in neotropic cormorants (*Phalacrocorax brasilianus*) in Chile.” Journal of Wildlife Diseases 55 (1): 105–112.

Verhagen, Josanne H., Frank Majoor, Pascal Lexmond, Oanh Vuong, Giny Kasemir, Date Lutterop, Albert D.M.E. Osterhaus, Ron A.M. Fouchier, and Thijs Kuiken. 2014. “Epidemiology of Influenza A Virus among black-headed Gulls, the Netherlands, 2006–2010.” Emerging Infectious Diseases 20, no. 1 (January): 138–141. issn: 1080-6059. 10.3201/eid2001.130984.

Wallensten, A., V. J. Munster, J. Elmberg, A. D. M. E. Osterhaus, R. A. M. Fouchier, and B. Olsen. 2005. “Multiple gene segment reassortment between Eurasian and American lineages of influenza A virus (H6N2) in guillemot *(Uria aalge)*.” Archives of Virology 150, no. 8 (May): 1685–1692. issn: 1432-8798.

Wille, Michelle, Yanyan Huang, Gregory J. Robertson, Pierre Ryan, Sabina I. Wilhelm, David Fifield, Alexander L. Bond, et al. 2014. “Evaluation of Seabirds in Newfoundland and Labrador, Canada, as Hosts of Influenza A Viruses.” Journal of Wildlife Diseases 50, no. 1 (January): 98–103. issn: 0090-3558. 10.7589/2012-10-247.

Wojczulanis-Jakubas, Katarzyna, Aleš Svoboda, Andrzej Kruszewicz, and Arild Johnsen. 2010. “No Evidence of Blood Parasites in Little Auks *(Alle alle)* Breeding on Svalbard.” Journal of Wildlife Diseases 46, no. 2 (April): 574–578. issn: 0090-3558. 10.7589/0090-3558-46.2.574.

Zagalska-Neubauer, Magdalena, and Staffan Bensch. 2015. “High prevalence of Leucocytozoon parasites in fresh water breeding gulls.” Journal of Ornithology 157, no. 2 (September): 525–532. issn: 2193-7206. 10.1007/s10336-015-1291-5.

## References

Battaglia, Michael, and Lee Ann Garrett-Sinha. 2023. “Staphylococcus xylosus and Staphylococcus aureus as commensals and pathogens on murine skin.” Laboratory Animal Research 39 (1): 18.

Blakeslee, April MH, Carolyn L Keogh, Amy E Fowler, and Blaine D Griffen. 2015. “Assessing the effects of trematode infection on invasive green crabs in eastern North America.” PLoS One 10 (6): e0128674.

Bock, Dieter, and Ottmar Janssen. 1987. “The life-cycle of Plagiorchis maculosus (Rudolphi, 1802) Braun, 1902 (Trematoda: Plagiorchiidae), a parasite of swallows (Hirundinidae).” Systematic parasitology 9 (3): 203–212.

Chai, Jong-Yil, and Bong-Kwang Jung. 2017. “Fishborne zoonotic heterophyid infections: an update.” Food and Waterborne Parasitology 8:33–63.

Cheong, Heng Choon, Chalystha Yie Qin Lee, Yi Ying Cheok, Grace Min Yi Tan, Chung Yeng Looi, and Won Fen Wong. 2019. “Chlamydiaceae: diseases in primary hosts and zoonosis.” Microorganisms 7 (5): 146.

Clifton, Ian J, and Daniel G Peckham. 2010. “Defining routes of airborne transmission of Pseudomonas aeruginosa in people with cystic fibrosis.” Expert Review of Respiratory Medicine 4 (4): 519–529.

Du, Xue-Yong, Pei-Fang Zhang, Sen-Rui Gong, Yuan-Sen Liang, Yu-Hao Huang, Hao-Sen Li, and Hong Pang. 2023. “Discovery of a novel circulation route of free-living Serratiasymbiotica mediated by predatory ladybird beetles.” FEMS Microbiology Ecology 99 (11): fiad133.

Galaktionov, KV, SWB Irwin, and DH Saville. 2006. “One of the most complex life-cycles among trematodes: a description of Parvatrema margaritense (Ching, 1982) n. comb.(Gymnophallidae) possessing parthenogenetic metacercariae.” Parasitology 132 (5): 733–746.

Gonchar, Anna, Damien Jouet, Karl Skírnisson, Darya Krupenko, and Kirill V Galaktionov. 2019. “Transatlantic discovery of Notocotylus atlanticus (Digenea: Notocotylidae) based on life cycle data.” Parasitology Research 118 (5): 1445–1456.

González-Acuña, Daniel A, Lucila Moreno, Michelle Wille, Bjorn Herrmann, Mike J Kinsella, and Ricardo L Palma. 2021. “Parasites of chinstrap penguins (*Pygoscelis antarctica*) from three localities in the Antarctic Peninsula and a review of their parasitic fauna.” Polar Biology 44 (11): 2099–2105.

Gray, Gregory C. 2006. “Adenovirus transmission—worthy of our attention.” The Journal of infectious diseases 194 (7): 871–873.

Hernández, AJ Chaves. 2014. “Poultry and avian diseases.” Encyclopedia of Agriculture and Food Systems, 504.

Hoberg, Eric P. 1986. “Evolution and historical biogeography of a parasite–host assemblage: Alcataenia spp.(Cyclophyllidea: Dilepididae) in Alcidae (Charadriiformes).” Canadian Journal of Zoology 64 (11): 2576–2589.

Lanao, Andrea E, Rebanta K Chakraborty, and Anthony L Pearson-Shaver. 2023. “Mycoplasma infections.” In StatPearls [Internet]. StatPearls Publishing.

Malik, Yashpal Singh, Arockiasamy Arun Prince Milton, Sandeep Ghatak, and Souvik Ghosh. 2022. “Avian Chlamydiosis (Psittacosis, Ornithosis).” In Role of Birds in Transmitting Zoonotic Pathogens, 137–147. Springer.

Martorelli, Sergio R, Brian L Fredensborg, Kim N Mouritsen, and Robert Poulin. 2004. “Description and proposed life cycle of Maritrema novaezealandensis n. sp.(Microphallidae) parasitic in red-billed gulls, Larus novaehollandiae scopulinus, from Otago Harbor, South Island, New Zealand.” Journal of Parasitology 90 (2): 272–277.

Murphy, John R. 1996. “Corynebacterium diphtheriae.” Medical Microbiology. 4th edition.

Rajapaksa, R, and CH Fernando. 1986. “Tropical species of Kurzia (Crustacea, Cladocera) with a description of Kurzia brevilabris sp. nov.” Canadian Journal of Zoology 64 (11): 2590–2602.

Sapiro, Anne L, Beth M Hayes, Regan F Volk, Jenny Y Zhang, Diane M Brooks, Calla Martyn, Atanas Radkov, Ziyi Zhao, Margie Kinnersley, Patrick R Secor, et al. 2023. “Longitudinal map of transcriptome changes in the Lyme pathogen Borrelia burgdorferi during tick-borne transmission.” Elife 12:RP86636.

Sewoyo, Palagan Senopati, I Putu Cahyadi Putra, and Willy Morris Nainggolan. 2024. “Pathology of proventricular tetrameriasis in a free-range chicken.” ARSHI Veterinary Letters 8 (3): 47–48.

Slifka, KM Jacobs, AE Newton, and BE Mahon. 2017. “Vibrio alginolyticus infections in the USA, 1988–2012.” Epidemiology & Infection 145 (7): 1491–1499.

Traversa, Donato, Angela Di Cesare, and Gary Conboy. 2010. “Canine and feline cardiopulmonary parasitic nematodes in Europe: emerging and underestimated.” Parasites & vectors 3 (1): 62.

Wong, PL, and RC Anderson. 1982. “The transmission and development of Cosmocephalus obvelatus (Nematoda: Acuarioidea) of gulls (Laridae).” Canadian Journal of Zoology 60 (6): 1426–1440.

Wozniakowski, Grzegorz, and Elżbieta Samorek-Salamonowicz. 2015. “Animal herpesviruses and their zoonotic potential for cross-species infection.” Annals of Agricultural and Environmental Medicine 22 (2).

Woźniakowski, Grzegorz, Maciej Frant, and Andrzej Mamczur. 2018. “Avian reticuloendotheliosis in chickens–an update on disease occurrence and clinical course.” Journal of Veterinary Research 62 (3): 257.

Zhao, Yang, Andre Aarnink, Maria Cambra-Lopez, and Teun Fabri. 2012. “Shedding and Emission of Airborne Viral Microorganisms from Animal Houses.” In 2012 IX International Livestock Environment Symposium (ILES IX), 3. American Society of Agricultural and Biological Engineers.

